# Metabolic basis of the astrocyte-synapse interaction governs dopaminergic-motor connection

**DOI:** 10.1101/2025.05.28.656515

**Authors:** Yanru Xu, Piaoping Kong, Mengqi Wang, Yanyun Mao, Zhiguo Ma

## Abstract

Perisynaptic astrocyte processes are constitutive attachments of synapses in the central nervous system. However, the molecular mechanisms that control perisynaptic astrocyte ensheathment and their implications in the wiring of neural circuits remain unclear. Here, we report that glycolysis controls astrocyte-synapse contact. In the *Drosophila* larval dopaminergic (DAergic) circuit, blocking astrocyte glycolysis stimulated perisynaptic ensheathment by attenuating astrocyte-to-DAergic neuron neuroligin 2-neurexin 1 signaling. As a result, the larvae executed more reorientation actions during locomotion. At the circuit level, behavioral alterations were found to arise from increased DAergic neuronal synaptogenesis and DAergic-motor connection. Our research uncovers an ancient metabolic basis that determines perisynaptic astrocyte ensheathment abundance through a conserved neuroligin-neurexin signaling pathway and demonstrates the role of astrocyte glycolysis in controlling DAergic-motor circuit assembly and function.

## Introduction

Astrocytes have a complex morphology with dense branches ensheathing many synapses^1–3^. Consequently, perisynaptic astrocyte processes and synapses are positioned in immediate vicinity for astrocyte-synapse interaction. Astrocyte-synapse contact is modulated by synaptic activity and several cell-cell communication pathways such as cell-adhesion signaling^4–14^.

Astrocytes are key non-neuronal cells that interact with the DAergic system^15–17^. The DAergic system comprises many subcircuits with patterned synaptic connectivity, each of which controls specific actions^18,19^. Imbalances in DAergic subcircuit connectivity are associated with both motor and non-motor deficits in neurodevelopmental disorders such as autism and schizophrenia^20–24^. Perisynaptic astrocyte processes regulate neurotransmission at DAergic synapses^25–28^; however, whether they contribute to patterning synaptic connection of developing DAergic subcircuits is unknown.

Neuroligins (Nlgs) and neurexins (Nrxs) are ligand-receptor, cell-adhesion molecules (CAMs) responsible for regulating synapse assembly and function^29–32^. The Nlg-Nrx signaling was traditionally viewed as a neuron-to-neuron communication pathway. For instance, postsynaptic Nlg2 and Nlg3 instruct DAergic presynapse organization^33,34^, and conditional knockouts of all Nrxs in DAergic neurons cause disproportionate co-transmission at DAergic synapses^35^. However, recent findings have provided new insights into the intercellular modalities of Nlg-Nrx signaling. Nonneuronal cells in the central nervous system (CNS), such as astrocytes, also express Nlgs^36,37^. In fact, astrocyte-neuronal Nlg-Nrx signaling is critical for excitation/inhibition balance and neural plasticity and regulates synaptogenesis in a circuit-specific manner ^8,38^. It is crucial to redefine the role of Nlg-Nrx signaling in the wiring of DAergic circuits; however, the question of how astrocyte-neuronal Nlg-Nrx signaling is regulated remains unclear.

Astocytes are efficient at glycolysis and supply the brain with glycolytic intermediates essential for neural biomass growth including synaptogenesis^39^, and the maintenance of neuronal activity^40^. Consistent with the established functions of vertebrate astrocytes, studies in *Drosophila* have demonstrated that astrocyte ablation impairs cholinergic synaptogenesis in the pupal antennal lobes^41^, while deficits in glial glycolysis lead to neuronal death in adult flies^42^. However, it remains to be elucidated whether the astrocyte-neuronal communication pathways, specifically Nlg-Nrx signaling and glycolysis, converge to regulate synaptogenesis and the precise wiring of neural circuit.

Here, using a previously unexplored *Drosophila* model addressing astrocyte-DAergic synapse interactions, we show the metabolic regulation of astrocyte-neuronal Nlg-Nrx signaling by glycolysis in controlling perisynaptic astrocyte ensheathment, DAergic-motor subcircuit wiring, and motor function. Our findings advance the understanding of astrocyte-synaptic mechanisms underlying the metabolic modulation of motor control by dopamine.

## Results

### Astrocyte ensheathment of DAergic synapses

First, to trace <u>p</u>erisynaptic <u>e</u>nsheathment by <u>a</u>strocytes in <u>p</u>roximity <u>o</u>f <u>D</u>Aergic <u>s</u>ynapses (hereafter referred to as PEAPODs), we expressed membrane-tethered split GFP (spGFP) fragments, CD4-spGFP^1–10^ and CD4-spGFP^11^, in astrocytes and DAergic neurons, respectively, in *Drosophila* larvae, using two binary expression systems, namely Gal4/UAS and LexA/lexAop (Fig. S1A)^15,43–46^. As several previous studies have modeled astrocyte interactions with the monoaminergic, motor, and GABAergic systems in the larval ventral nerve cord (VNC), we also chose to focus on the VNC^15,38,41^. Using GFP reconstitution, we identified many astrocyte-DAergic neuronal contact sites, which disperse along DAergic neuronal processes and show regional variations in intensity across the VNC and brain (Fig. S1B). Many of them resembled synaptic boutons and adjoined or contained core active zone protein Bruchpilot (Brp), indicative of perisynaptic astrocyte ensheathment of DAergic synapses (i.e., PEAPODs) (Fig. 1A; Supplemental video S1). This differs from larval glutamatergic circuit where astrocyte-synapse contact is rare^47^. To quantitate the astrocyte-DAergic synapse association, we examined PEAPODs and DAergic neurons expressing the presynaptic marker Bruchpilot^short^-mStraw (Brp.S-mStraw) at single-cell resolution using FLP-out^44,48^. As a result, we found that 87.5% ± 0.55% of Brp.S-mStraw puncta were associated with PEAPODs in the VNC, and the number dropped to 41.9% ± 1.07% in the brain (Figs. 1B–1D; Fig. S1C).

**Figure 1.**
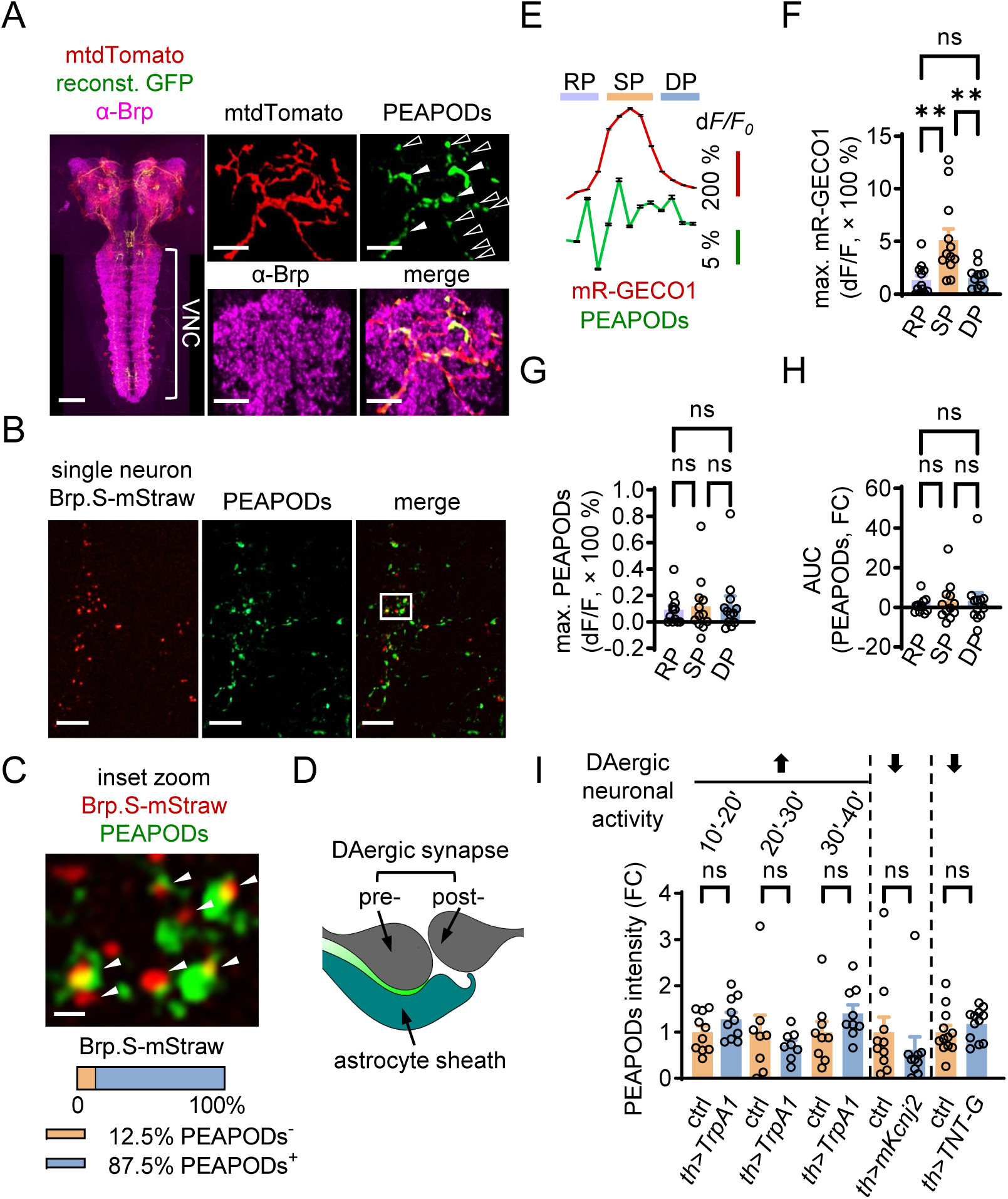
Perisynaptic astrocyte ensheathment of DAergic synapses. (A) Third instar larval CNS with the VNC highlighted (scale bar: 50 µm). Reconstituted GFP (reconst. GFP) reveals astrocyte contact sites (arrowheads and open arrowheads) with DAergic neuronal processes labeled by myristoylated tdTomato (mtdTomato) (scale bars: 5 µm). Open arrowheads identify typical PEAPODs that resemble synaptic boutons on terminal branches of DAergic neurons. Brp: Bruchpilot (presynaptic marker). (B) Single-cell PEAPODs and synaptic labeling of the DAergic neuron in larval ventral nerve cord (n = 11) (scale bars: 10 µm). (C) Inset zoom in (B), arrowheads denote PEAPODs^+^ Brp.S-mStraw puncta (scale bar: 2 µm). (D) Schematic drawing of the PEAPODs. Note that GFP reconstitutes at the contact site between the astrocyte sheath and DAergic synapse. (E–H) PEAPODs intensity vs. spontaneous DAergic neuronal burst activation (Ca^2+^, mR-GECO1) (n = 12). RP, SP, and DP: rising, sustaining, and decay periods, respectively. The area under the curve (AUC) was calculated to show the accumulative PEAPODs response over time. max.: Maximum. (I) PEAPODs intensity. Genetically manipulated DAergic neuronal activity (arrows, up or down) as indicated (n = 8–12). *th*: DAergic neuron driver. *mKcnj2*, *TNT-G* inactive controls: *mKcnj2.nc*, *TNT-Q*. (F), (G), (H), one-way ANOVA with Tukey’s test. (I), unpaired t-test. \*\**P* < 0.01. ns: Not significant, FC: Fold change.

Neuronal activity remodels astrocyte-synapse contact, and DAergic neurons spontaneously fire in bursts^15,49^. To determine how PEAPODs reorganize during DAergic neuronal activation, we performed dual-color live imaging of PEAPODs and measured the activity of DAergic neurons using the membrane-tethered Ca^2+^ indicator mR-GECO1 (Fig. S1D; Supplemental video S2)^50^. We recorded active DAergic neuronal branches that appeared to be transversal projections from vmTH neurons^51^. Spontaneous DAergic neuronal bursting plotted a bell-shaped Ca^2+^ curve, for which we named three phases: rising, sustaining, and decay periods (RP, SP, and DP, respectively) (Fig. 1E). The RP and DP Ca^2+^ levels (135.2% ± 12.21% vs. 152.1% ± 9.71%, maximum normalized difference in fluorescence levels between any given time point and the basal state [dF/F]) were similar and dramatically lower than those of the SP (512.1% ± 31.21%, maximum dF/F) (Fig. 1F). PEAPODs presented a multiple-peak response during DAergic neuronal burst firing (Fig. 1E). In contrast to the bell-shaped DAergic neuronal Ca^2+^ activity with a pair of symmetrical RP, DP shoulders, and an SP peak, the PEAPODs exhibited equivalent increases in intensity during the RP, SP, and DP (9.2% ± 0.97%, 11.9% ± 1.85%, and 12.9% ± 1.94%, respectively; maximum dF/F) (Figs. 1G and 1H), before returning to baseline. Therefore, spontaneous DAergic neuronal activity is not sufficient to stabilize temporarily enhanced PEAPODs. Thermogenetic activation of DAergic neurons for 30 min using the cation channel TrpA1 did not change the intensity of the PEAPODs (Fig. 1I). Finally, we inhibited DAergic neurons by hyperpolarizing the membrane potential using the potassium inwardly rectifying channel mKcnj2 or by blocking synaptic transmission with the tetanus toxin TNT-G, with none showing significantly altered PEAPODs intensity (Fig. 1I). Taken together, these results suggest that astrocyte-DAergic synapse contact is transiently stimulated during spontaneous DAergic neuronal burst firing, but that DAergic neuronal activity is dispensable for PEAPODs.

### Astrocytes determine the abundance of PEAPODs via glycolysis

Glycolysis breaks down one glucose molecule into two molecules of pyruvate (Fig. 2A). Vertebrate astrocytes exhibit high glycolytic activity^52,53^, and glial glycolysis has been shown critical for neuronal survival in adult flies^42^. We found that the knockdown (KD) of glycolytic genes by RNAi, including *phosphofructokinase* (*pfk*), *enolase* (*eno*), and *pyruvate kinase* (*pyk*), in astrocytes increased the intensity and number of PEAPODs by several folds: *pfk* RNAi (e.g., 5.6 ± 0.12, 2.2 ± 0.02), *eno* RNAi (e.g., 7.0 ± 0.18, 2.4 ± 0.05), and *pyk* RNAi (e.g., 5.8 ± 0.10, 2.0 ± 0.02) (intensity, number; fold change). In addition, knocking down these genes in astrocytes caused many PEAPODs to enlarge hundreds of times in volume (Figs. 2B–2E, S2A–S2D).

**Figure 2.**
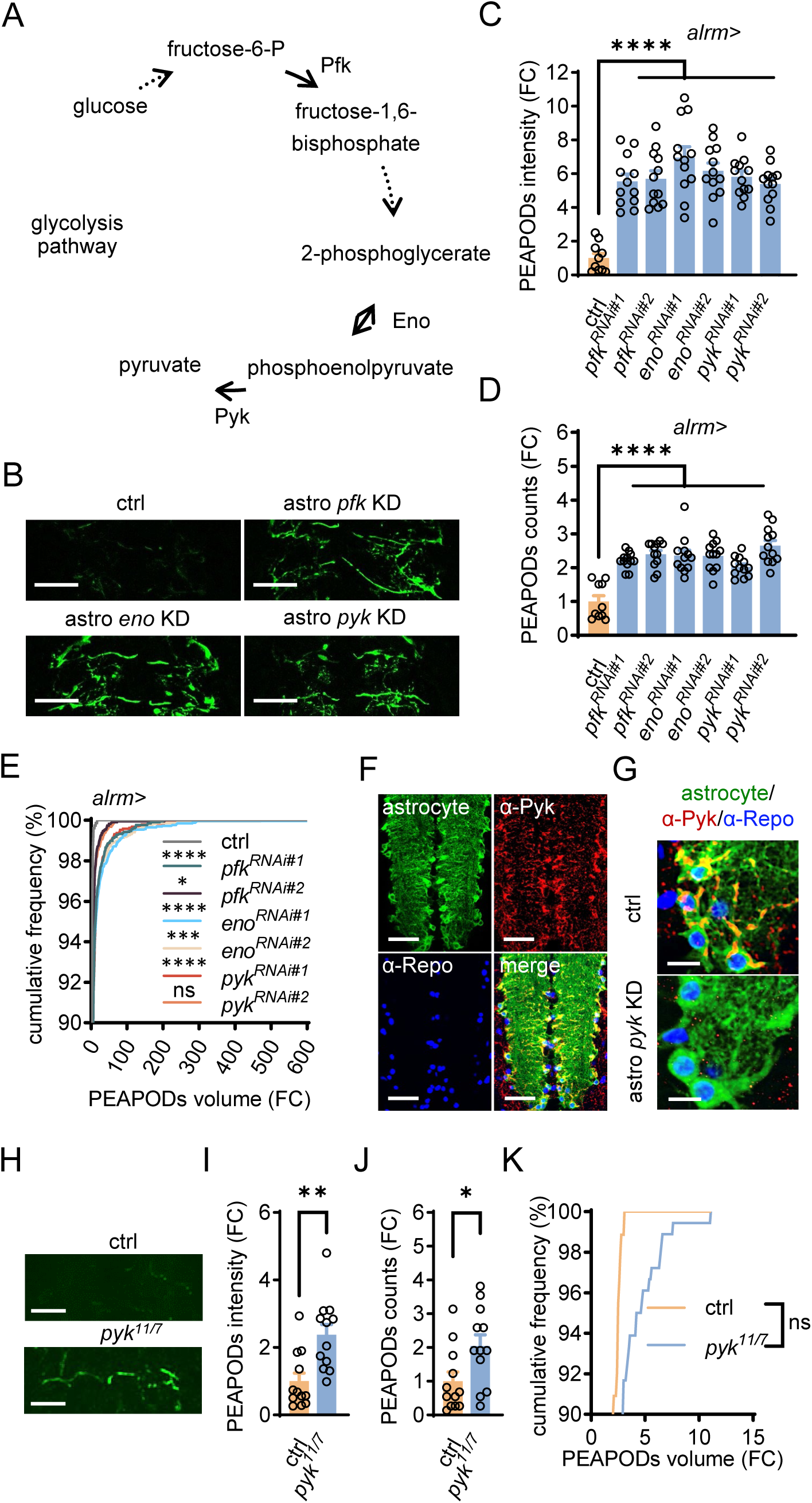
Astrocyte glycolysis deficiency increases PEAPODs abundance. (A) Biochemical pathway of glycolysis. Dotted arrows indicate the intermediate steps not shown. The double-headed arrow denotes reversible reactions. (B–E) PEAPODs and abundance analysis (scale bars: 50 µm). Gene knockdown (KD) in the astrocyte (astro) using RNAi were driven by the *alrm* driver. Genotypes as indicated (n = 10–12. In E, 784–2583 individual PEAPODs in total). (F) Pyk (α-Pyk) was enriched in astrocytes (scale bars: 50 µm). (G) *pyk* KD diminished Pyk expression (scale bars: 10 µm). In (F) and (G), mCD8-GFP labeled astrocytes, and α-Repo stained glial cell nuclei. (H–K) PEAPODs and abundance analysis (scale bars: 25 µm). Genotypes as indicated (n = 10. In K, 88–180 individual PEAPODs in total). (C), (D), one-way ANOVA with Tukey’s test. (I), (J), unpaired t-test. (E), (K), Kolmogorov-Smirnov test. \**P* < 0.05, \*\**P* < 0.01, \*\*\**P* < 0.001, \*\*\*\**P* < 0.0001. ns: Not significant, FC: Fold change.

Pyk is the final-step, rate-limiting enzyme that converts phosphoenolpyruvate to pyruvate during glycolysis. Anti-Pyk antibody staining showed that Pyk was enriched in astrocytes (Fig. 2F), and astrocyte *pyk* KD diminished its expression (Fig. 2G). Next, we recovered two *pyk* alleles by CRISPR-Cas9 mutagenesis^54–57^, *pyk*^11^ and *pyk*^7^, which carried 1-bp (*pyk*^11^) and 17-bp (*pyk*^7^) frameshift deletions in the largest *pyk* exon, respectively (Fig. S3A). Pyk expression was severely reduced in the mutants (e.g., 27.7% ± 4.68% remained in *pyk*^11^*^/^*^7^) (Figs. S3B and S3C), which caused the mutants to die at the 2^nd^ instar larval stage. Similar to astrocyte *pyk* RNAi, *pyk* mutations also increased PEAPODs (Figs. 2H–2K). In conclusion, astrocytes determine the PEAPODs abundance by glycolysis.

### Glycolysis-deficient astrocytes promote DAergic neuron synaptogenesis

*Drosophila* astrocytes exhibit a striking homology with their vertebrate counterparts in morphology and function. For example, astrocytes, characteristic of sophisticated bushy processes in both species, infiltrate the neuropile where they ensheath synapses and play a key role in synaptogenesis^41,58–65^. We labeled astrocytes with membrane-tethered GFP (i.e., mCD8-GFP) and observed equivalent amounts of astrocytes and normal neuropile infiltration in the *pyk* mutant compared to the control (Figs. 3A and 3B). Using anti-Brp antibody staining, we detected similar levels of Brp in the *pyk* mutant compared to the control, suggesting that overall CNS synaptogenesis is not affected by glycolysis deficiency (Fig. 3C). To examine astrocyte morphology, we conducted mosaic analysis with a repressible cell marker (MARCM) to label single astrocytes^66^. We found *pyk* mutant astrocytes formed thick branchlets and radiated out to the size of control (Figs. 3D and 3E).

**Figure 3.**
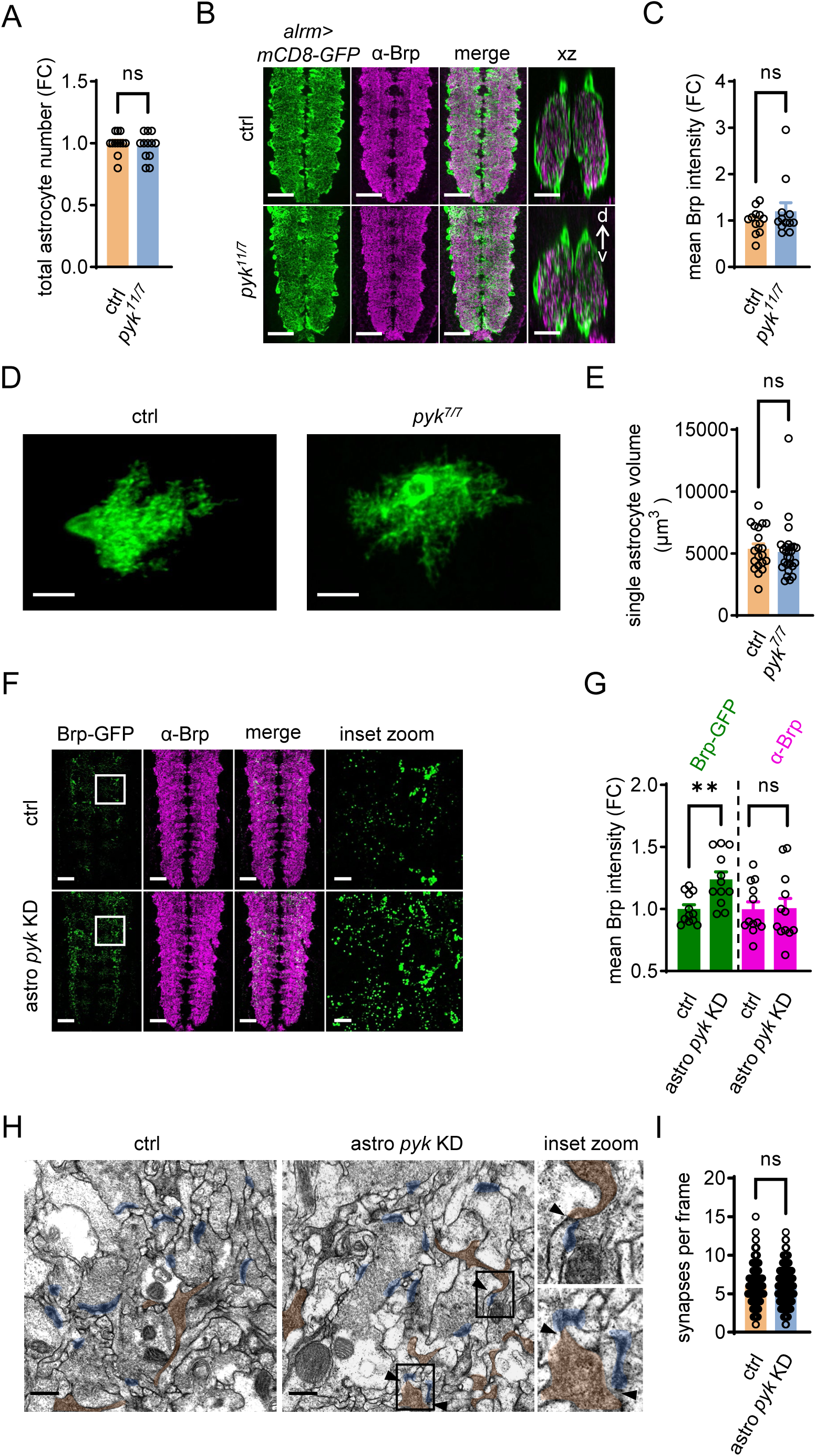
Glycolysis-deficient astrocytes promote DAergic synaptogenesis. (A) Astrocyte number. Genotypes as indicated (n = 12). (B) Normal neuropile (marked by α-Brp) infiltration by the astrocyte in the *pyk*^11^*^/^*^7^ mutant (scale bars: 20 µm). xz confocal sections show the infiltration in the dorsal (d)-ventral (v) axis of the VNC. Genotypes as indicated (n = 12). *alrm*: Astrocyte driver, mCD8-GFP: Membrane-tethered GFP, Brp: Bruchpilot (presynaptic marker). (C) Mean Brp levels. Genotypes as indicated (n = 12). (D and E) Single astrocyte MARCM clone. Genotypes as indicated (n = 20–26). MARCM: mosaic analysis with a repressible cell marker. (F and G) Endogenous Brp (Brp-GFP) in DAergic neurons (scale bars: 25 µm; 5 µm in the inset zoom). Genotypes as indicated (n = 12). astro: Astrocyte, KD: Knockdown. (H and I) Sections by transmission electron microscopy (scale bars: 500 nm). Color shade: synapses, blue; astrocyte processes, orange. Note that perisynaptic astrocyte process (arrowheads) were more often observed in astro *pyk* KD. Genotypes as indicated (n = 311–313 frames from 5 larval CNS). Frame size, 4.28 μm (width) × 4.41 μm (length). astro: Astrocyte, KD: Knockdown. Unpaired t-test. \*\**P* < 0.01. ns: Not significant, FC: Fold change.

We showed that astrocyte glycolysis deficiency increased the abundance of PEAPODs. Therefore, we next examined whether DAergic synaptogenesis was altered in turn. To this end, we specifically labeled endogenous Brp in DAergic neurons with a flip-out cassette, *Brp>FSF>GFP*, using STaR^67^. We created a *lexAop2-pyk^RNAi^* transgenic line to knock down *pyk* in astrocytes (Fig. S3D). Astrocyte *pyk* KD using the LexA/lexAop system showed identical PEAPODs enhancement phenotypes (Figs. S3E–S3G). We observed that astrocyte *pyk* KD increased endogenous Brp expression in DAergic neurons to 123.8% ± 1.75% of the control (Figs. 3F and 3G), while the overall CNS synaptogenesis remained unchanged (α-Brp and transmission electron microscopy; Figs. 3H and 3I). Thus, astrocyte glycolysis deficiency promotes DAergic neuron synaptogenesis. However, we do not rule out that non-DAergic synaptogenesis is also altered by astrocyte glycolysis deficiency. It is unlikely that Brp stimulates PEAPODs because the *brp* KD in DAergic neurons enhances PEAPODs (Figs. S4A–S4C).

### Astrocyte glycolysis deficiency leads to biased DAergic circuit wiring

We demonstrated that astrocyte glycolysis deficiency increased PEAPODs and DAergic neuron synaptogenesis. However, whether this had a significant effect on DAergic circuit assembly remains an open question. In *Drosophila*, postsynaptic connections with the DAergic system remain largely unknown, particularly those in the larval VNC. To address this issue, we profiled postsynaptic partners wired to DAergic neurons in the larval CNS. Briefly, we labeled the postsynaptic cells with mtdTomato by transsynaptic tracing (i.e., *trans*-Tango) and pooled mtdTomato^+^ cells using fluorescence-activated cell sorting (FACS)^68,69^. Subsequently, we performed single-cell RNA sequencing (scRNA-seq) and cell type annotation to identify the cohort-based postsynaptic connectome of the larval DAergic system (Fig. 4A).

**Figure 4.**
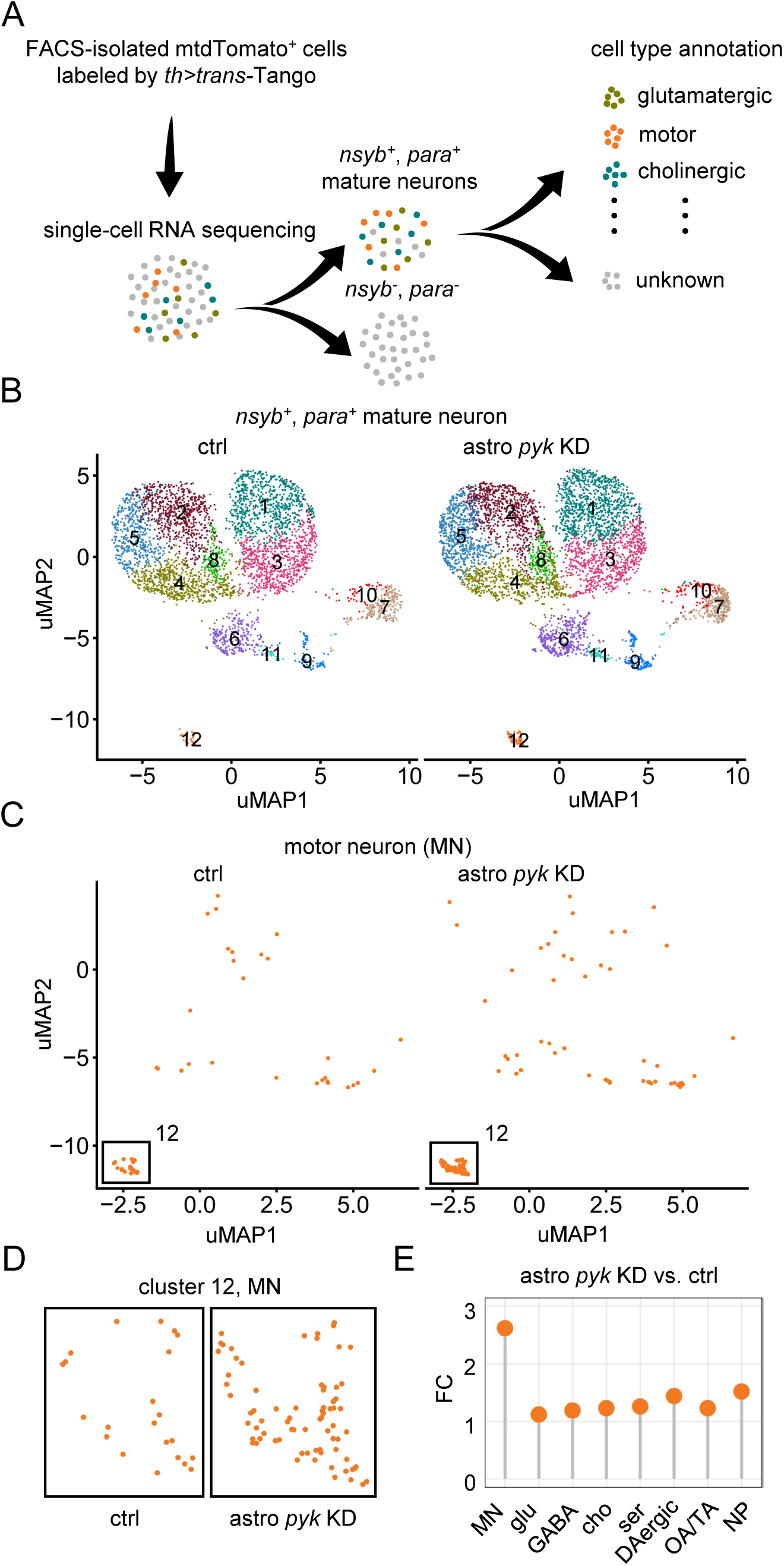
Astrocyte glycolysis deficiency causes biased DAergic wiring. (A) Schematic protocol for profiling postsynaptic partners (labeled by *trans*-Tango) wired to DAergic neurons. FACS: Fluorescence-activated cell sorting, *th*: DAergic neuron driver, *trans*-Tango: Transsynaptic labeling system. (B) Clustering of *nsyb^+^*, *para^+^* mature neurons (ctrl, n = 5589; astrocyte [astro] *pyk* knockdown [KD], n = 7236). Note that cluster 12 is distant to the others. uMAP: Uniform manifold approximation and projection. (C) Annotated MNs (ctrl, n = 50; astro *pyk* KD, n = 131) in the UMAP. MNs gathered in cluster 12. (D) Close-up view of cluster 12 MNs. (E) Plotting FCs in the cell number of annotated populations, astro *pyk* KD vs. ctrl. The MN showed the highest FC and followed the neuropeptidergic (NP) neuron and the DAergic neuron. glu: Glutamatergic, GABA: GABAergic, cho: Cholinergic, ser: Serotoninergic, OA/TA: Octop-/tyr-aminergic, FC: Fold change.

We dissected 50 CNSs each from the control and astrocyte *pyk* KD larvae, ultimately obtaining 11815 and 13327 (control vs. astrocyte *pyk* KD) mtdTomato^+^ cells, respectively, which were sequenced. The higher yield of mtdTomato^+^ cells from astrocyte *pyk* KD is in line with our observation that KD led to stronger mtdTomato expression by *trans*-Tango, which suggests that DAergic neurons make more postsynaptic connections (Figs. S4D and S4E). These individual cells were grouped into 20 clusters in uniform manifold approximation and projection (uMAP) analysis (Fig. S5A). *nsyb^+^para^+^* cells were selected as mature neurons^70,71^, which accounted for 47.3% (control) and 54.3% (astrocyte *pyk* KD) of all cells sequenced (Figs. S5B–S5E). The mature neurons re-clustered in 12 groups (Fig. 4B). Next, using unique marker genes^71,72^, we performed cell type annotation and defined eight mature neuron populations (Figs. S5F, S5G, and S6A–S6J; Table S1)^73^. Our annotation is of high accuracy: for instance, 93.9% *twit*^+^ motor neurons (MNs) can be sorted into known MN subclasses with an additional marker gene, e.g., 53.6% glutamatergic type I (*vglut*^+^), 18.2% octopaminergic type II (*tβh*^+^), and 22.1% neuropeptidergic type III (*phm*^+^) (Table S2)^74,75^. Moreover, excitatory/inhibitory populations appeared to be mutually exclusive in the uMAP analysis, e.g., glutamatergic vs. GABAergic populations (Figs. S6B vs. S6C) and GABAergic vs. cholinergic populations (Figs. S6C vs. S6D). We compared the cell numbers of each population in the control and astrocyte *pyk* KD, with the astrocyte *pyk* KD often outnumbering the controls. The MNs, gathered in cluster 12, had the highest fold change (>2.6), followed by the neuropeptidergic neurons (>1.5) and DAergic neurons (>1.4) (Figs. 4C–4E). Therefore, enhanced DAergic synaptogenesis caused by astrocyte glycolysis deficiency indeed gives rise to more postsynaptic connections. Taken together, our data demonstrate that astrocyte glycolysis deficiency leads to biased DAergic circuit wiring.

### Biased DAergic-motor wiring alters motor output

Dopamine modulates motor circuit dynamics^76–80^. To examine whether biased DAergic-motor wiring alters motor output, we tracked and analyzed *Drosophila* larval locomotion using the frustrated total internal reflection (FTIR)-based imaging method^81,82^. This larval behavior manifests as a series of active forward, as well as a few backward, crawling (i.e., peristalsis) actions interrupted by stereotyped reorientation attempts (i.e., pauses and sweeps) (Fig. 5A)^83–85^. Compared to the controls, the astrocyte *pyk* KD larvae travelled in densely branched trajectories (Fig. 5B). They increased the likelihood of pausing (e.g., 16.4% ± 0.22% vs. 6.0% ± 0.27%, astrocyte *pyk* KD vs. ctrl) and sweeping (e.g., 16.8% ± 0.23% vs. 5.9% ± 0.26%, astrocyte *pyk* KD vs. ctrl) without changing bending strength (Figs. 5C–5E). During the peristaltic sessions, the RNAi larvae attained a higher maximum speed (e.g., 2.7 mm/s ± 0.04 mm/s vs. 1.9 mm/s ± 0.02 mm/s, astrocyte *pyk* KD vs. ctrl) and maintained similar peristalsis efficiency and frequency (Figs. 5F–5H).

**Figure 5.**
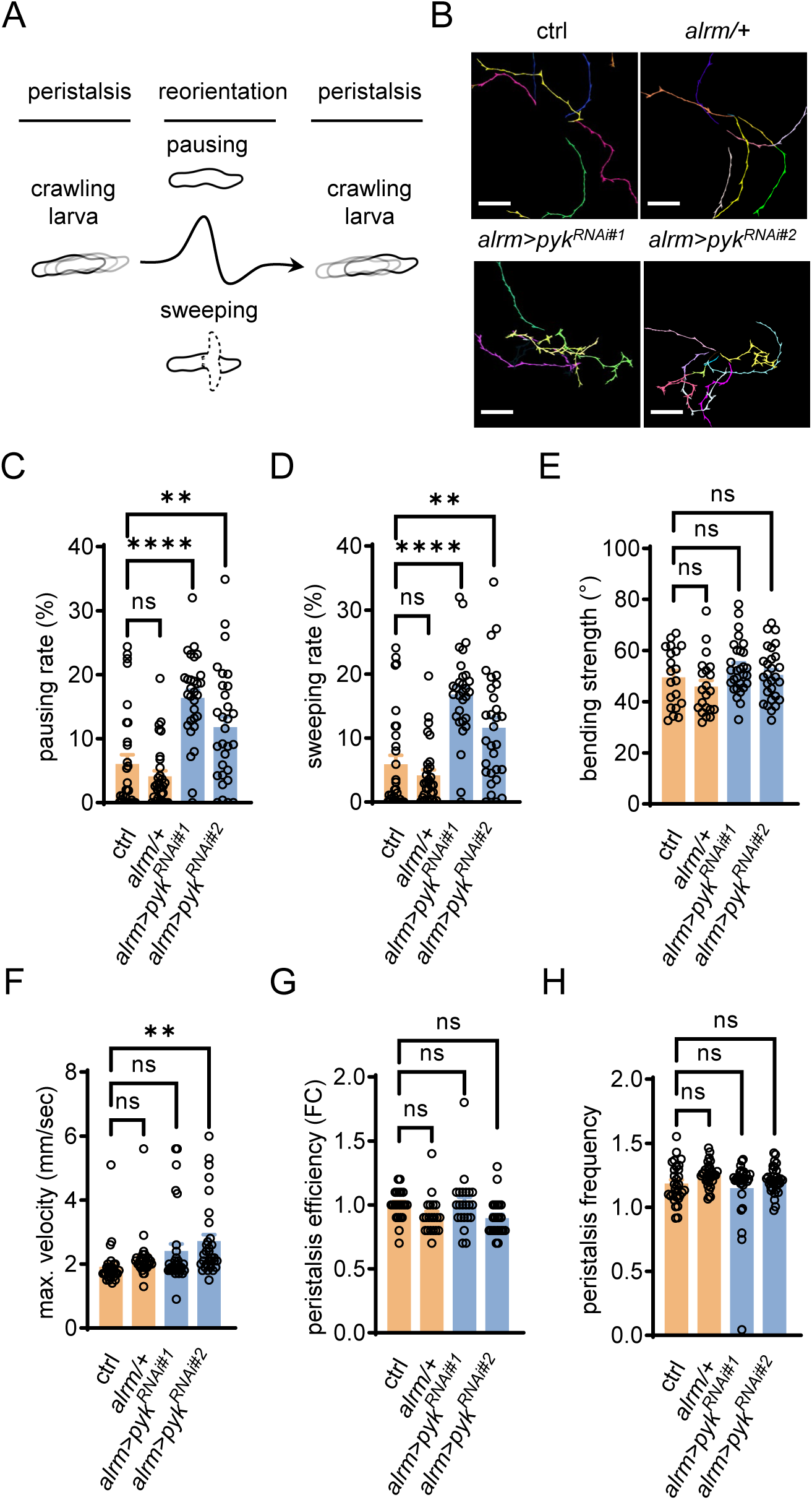
Biased DAergic wiring alters locomotion patterns. (A) Schematic illustration of *Drosophila* larval locomotive movements. Reorientation attempts intervene between peristaltic sessions. (B) Recorded larval locomotion trajectories (scale bars: 2 cm). Astrocyte *pyk* RNAi larvae travelled in paths with dense branches; six larvae per recording. Genotypes as indicated (n = 30). *alrm*: Astrocyte driver. (C–H) Quantitative analysis of larval locomotion. Genotypes as indicated (n = 30). Pausing rate (C): The number of frames the larva pauses/the total number of frames the larva appears × 100%, sweeping rate (D): The number of frames the larva sweeps/the total number of frames the larva appears × 100%, bending strength (E): Median bending angle, max. velocity (F): Maximum velocity, peristalsis efficiency (G): Distance per peristalsis, peristalsis frequency (H): Peristalses per second). One-way ANOVA with Tukey’s test. \*\**P* < 0.01, \*\*\*\**P* < 0.0001. ns: Not significant, FC: Fold change.

### Glycolysis controls the astrocyte-DAergic synapse interaction by Nlg2-Nrx1 signaling

It remains to be determined how astrocytes transduce the glycolytic event into a means of cell-cell communication guiding astrocyte-DAergic synapse interaction. CAMs such as Nlgs and Nrxs modulate astrocyte-synapse contact and synaptogenesis^8,38^. We assayed the effect of KDs of Nlgs and Nrxs on PEAPODs by RNAi and found that both astrocyte *nlg2* KD and DAergic neuron *nrx1* KD increased PEAPODs, e.g., astrocyte *nlg2* RNAi (5.2 ± 0.08, 2.9 ± 0.04) and DAergic neuron *nrx1* RNAi (2.4 ± 0.04, 1.9 ± 0.03) (intensity, number; fold change) (Figs. 6A–6H). In addition, *nrx1* RNAi in DAergic neurons enhanced synaptogenesis to 112.8% ± 0.63% of the control (Figs. 6I–6J). In summary, these data suggest that astrocyte glycolysis and astrocyte-DAergic neuronal Nlg2-Nrx1 signaling are somehow in the same pathway responsible for controlling PEAPODs and DAergic synaptogenesis. Using anti-Nlg2 antibody staining, we observed massive intra-astrocyte membranous Nlg2 bodies in the *pyk* mutant (Figs. 6K and 6L). The Nlg2 bodies were decorated with the endoplasmic reticulum (ER) luminal reporter GFP-KDEL, indicating that Nlg2 ER exit fails in *pyk* mutant astrocytes. Taken together, our findings support the notion that astrocyte glycolysis enables astrocyte-to-DAergic neuron Nlg2-Nrx1 signaling to restrict astrocyte-DAergic synapse contact and DAergic synaptogenesis.

**Figure 6.**
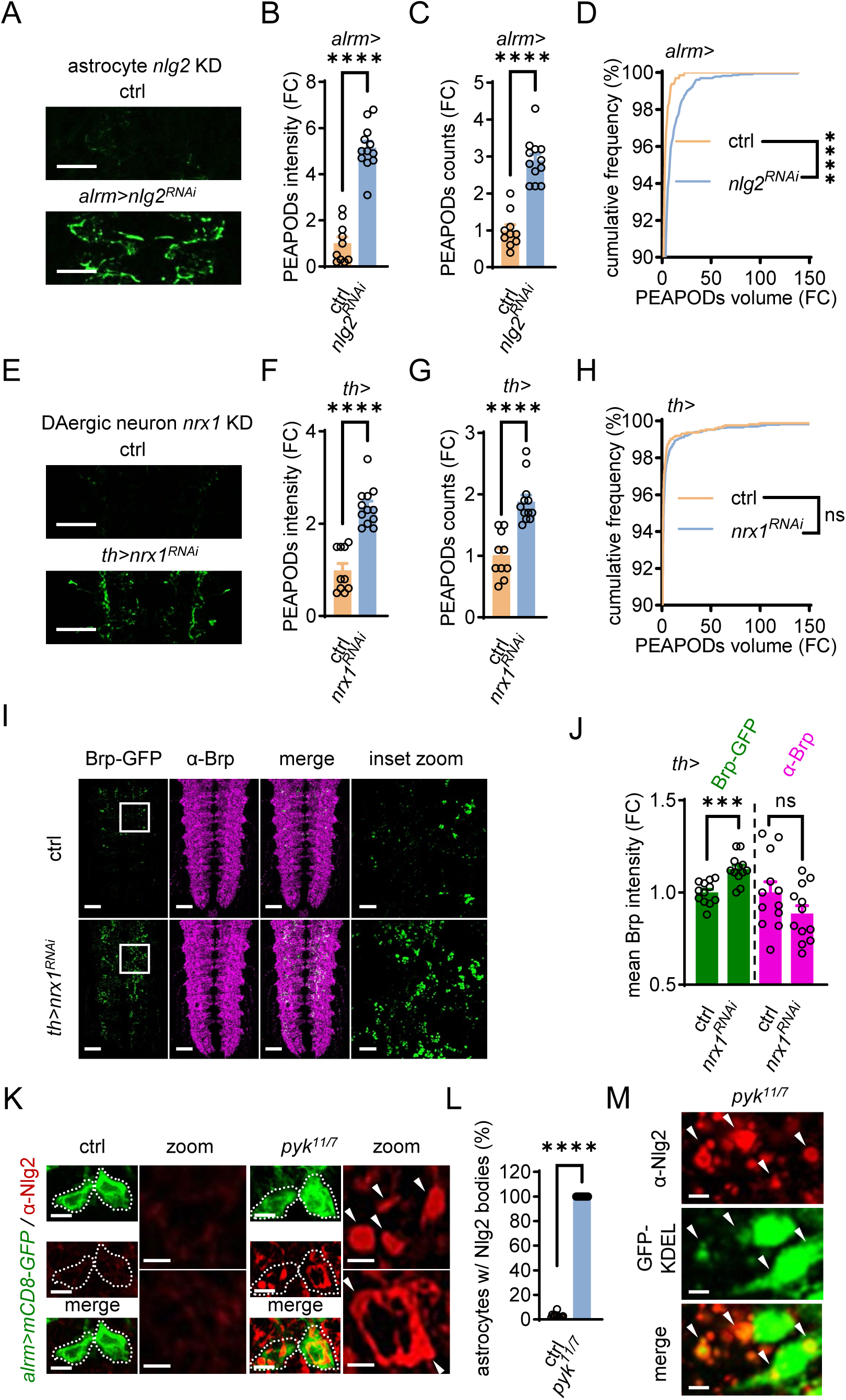
Astrocyte glycolysis deficiency disables astrocyte-DAergic neuronal Nlg2-Nrx1 signaling. (A–H) PEAPODs and abundance analysis (scale bars: 50 µm). Genotypes as indicated (n = 10–12; in D, 659–2286; in H, 1756–3975 individual PEAPODs in total). *alrm*: Astrocyte driver, *th*: DAergic neuron driver, KD: Knockdown. (I and J) Endogenous Brp (Brp-GFP) in DAergic neurons (scale bars: 25 µm; 5 µm in the inset zoom). Genotypes as indicated (n = 12). Brp: Bruchpilot (presynaptic marker). (K and L) Glycolysis deficiency led to intra-astrocyte Nlg2 bodies (arrowheads) (scale bars: 5 µm; 2 µm in the inset zoom). The dotted line marks the astrocyte soma. Genotypes as indicated (n = 12). mCD8-GFP: Membrane-tethered GFP. (M) The ER reporter GFP-KDEL decorated Nlg2 bodies (arrowheads) (scale bars: 2 µm). (B), (C), (F), (G), (J), (L), unpaired t-test. (D), (H), Kolmogorov-Smirnov test. \*\*\**P* < 0.001, \*\*\*\**P* < 0.0001. ns: Not significant, FC: Fold change.

## Discussion

Our data indicate a metabolic basis for perisynaptic astrocyte ensheathment. Glycolysis is critical for proper Nlg2 trafficking in astrocytes, and astrocyte-DAergic neuronal Nlg2-Nrx1 signaling relies on astrocyte glycolysis. This metabolic signaling axis (glycolysis/Nlg2-Nrx1) restricts perisynaptic astrocyte ensheathment; once it fails during development, the astrocyte-synapse contact rises and DAergic neurons increase synaptogenesis and their synaptic connection with motor neurons.

Nlg family proteins are transported to the cell surface where they bind Nrx receptors^86^. Nlgs contain many amino acid residues involved in N- and O-linked glycosylation^87,88^. Posttranslational glycosylation is crucial for Nlgs on their way to activating Nrxs, from protein folding to surface trafficking, as well as Nrx binding^89,90^. Autism-associated mutations of several *nlg* genes lead to differentially glycosylated Nlg proteins and their retention at the ER^91–93^. In this study, we showed that Nlg2 (mammalian Nlg1 homolog) is sequestered at the ER in glycolysis-deficient astrocytes. Modulations in glycolytic flux alter protein glycosylation outcomes^94–99^. Taken together, our findings suggest a similar mechanism for Nlg2 retention in the ER owing to aberrant glycosylation in glycolysis-deficient astrocytes. As a result, perisynaptic astrocyte ensheathment increases at DAergic synapses.

In terms of how perisynaptic astrocyte ensheathment abundance is relevant to synaptogenesis, previous studies have shown that astrocyte Nlg2 knockout reduced both perisynaptic astrocyte contact and glutamatergic synaptogenesis^8^. Additionally, the depletion of neuronal cell adhesion molecule (NRCAM) in astrocytes enhances astrocyte-synapse adhesion but decreases GABAergic synaptogenesis^10^. In contrast, our data indicate that astrocyte Nlg2 deficiency increases both perisynaptic astrocyte ensheathment and DAergic synaptogenesis. In summary, these findings suggest that perisynaptic astrocyte ensheathment abundance is a proxy for measuring astrocyte-synapse interaction but is not directly or inversely proportionally linked to synaptogenesis. Complex contact-mediated astrocyte-synaptic mechanisms regulate synaptogenesis, which involve the heterogeneity of synapses and neural circuits.

We mapped cohort-based postsynaptic connectomes of the *Drosophila* larval DAergic system using transsynaptic tracing, scRNA-seq, and cell type annotation. The connectomes comprise several heterogeneous populations. Most of the populations were annotated using defined marker genes associated with neurotransmitter types but not behavioral signatures. Therefore, our current analyses have a limited depth with regard to analyzing DAergic subcircuits associated with specific behaviors. However, the connectomes we constructed are valuable reference landscapes for DAergic subcircuit mapping when unique marker genes emerge in the midst of investigating dopamine signaling during *Drosophila* development. For instance, in this study, we observed a single molecularly defined population, the MNs, designated for locomotion. We found that astrocyte glycolysis deficiency led to more than double the number of MNs in synaptic connection with the DAergic system. Notably, astrocyte Nlg2 depletion extends the critical window of *Drosophila* larval motor circuit, which induces long-term changes in motor circuit wiring and behavior during development^38^. Similarly, we showed that astrocyte glycolysis deficiency causes astrocyte-neuronal Nlg2-Nrx1 signaling to fail and enhances DAergic synaptogenesis. Taken together, the data suggest that *Drosophila* astrocytes control the wiring of DAergic circuit among others by glycolysis. DAergic neurons start to appear as early as embryogenesis stage 14, before the motor critical window closes normally, beginning a few hours after larval hatching^38,100^. It remains unclear whether larval motor circuit plasticity plays a role in favoring DAergic-motor connection due to astrocyte glycolysis deficiency.

Primitive glycolysis is a fundamental metabolic process in astrocytes^101^. Our study opens new opportunities for uncovering the neurometabolic implications in neural circuit connectivity, particularly of the DAergic system.

## Author Contributions

Conceptualization: Z.G.M.; Methodology: Z.G.M., Y.R.X.; Investigation: Z.G.M., Y.R.X., P.P.K., M.Q.W., Y.Y.M.; Validation: Z.G.M., Y.R.X., M.Q.W.; Visualization: Z.G.M., Y.R.X.; Formal analysis: Z.G.M., Y.R.X.; Data curation: Z.G.M., Y.R.X.; Resources: Z.G.M.; Software: Z.G.M.; Project administration: Z.G.M.; Funding acquisition: Z.G.M.; Supervision: Z.G.M.; Writing – original draft: Z.G.M., Y.R.X.; Writing – review & editing: Z.G.M.

## Acknowledgements

This work was supported by National Excellent Young Scientists Fund (Overseas) and in part by Non-profit Central Research Institute Fund of Chinese Academy of Medical Sciences (2023-PT310-01). We thank the core facilities of Liangzhu Laboratory and the core facilities of Zhejiang University School of Medicine for the support in confocal microscopy and transmission electron microscopy. We acquired many valuable fly strains from Bloomington *Drosophila* Stock Center, TsingHua Fly Center and Vienna *Drosophila* Resource Center, we are grateful for their efforts.

## Materials and Methods

### *Drosophila* husbandry and stocks

Flies were cultured in cornmeal food at 25°C, unless stated otherwise. Detailed stock information is provided in Table S4.

### Molecular biology and transgenic flies

The construct *lexAop2-pyk^RNAi^* was assembled in alignment with attP/attB site-directed transgene integration in *Drosophila*, as illustrated in Fig. S3D. The *lexAop2-pyk^RNAi^* transgenic fly was obtained via standard germline microinjection.

### Single-cell clone induction

The genotype of larvae for single-neuron PEAPODs and synaptic labeling: *alrm-LexA, lexAop-CD4-spGFP*^11^*/UAS-brp.S-mStraw*; *hs-FLPG5, tub>Gal80>/th-Gal4, UAS-CD4-spGFP*^1–10^. Eleven single VNC DAergic neurons (out of 120 3^rd^ instar larval CNSs dissected) were obtained by 37°C heatshock for 10 min at 22 h after egg laying. Confocal images were taken using a Zeiss LSM 900 with Airyscan (objective lens, Plan-Apochromat 63×/1.40 NA, Oil, DIC M27). To quantitate astrocyte-DAergic synapse association: Image J plugin 3D objects counter was used to create two new image stacks by segmenting Brp.S-mStraw puncta and PEAPODs in confocal images from separate channels. The two image stacks (Brp.S-mStraw objects [BO], PEAPODs objects [PO]) were merged, and the association between BO and PO was examined.

Genotypes for generating clones by mosaic analysis with a repressible cell marker (MARCM): control, *alrm-Gal4, UAS-mCD8-GFP/repo-FLP; FRT82B/FRT82B, tub-Gal80*; *pyk* mutant, *alrm-Gal4, UAS-mCD8-GFP/repo-FLP; FRT82B, pyk*^7^*/FRT82B, tub-Gal80*. We took confocal images with an inverted Olympus IXplore SpinSR spinning disk confocal super resolution microscope (objective lens, UPLXAPO 60×/1.5 NA, Oil) and measured astrocyte volume in Imaris software.

### Live imaging and data analysis

We followed the live imaging protocol described by Ma et al. with minor modifications^1^. The 3^rd^ instar larval CNS was dissected in low Ca^2+^ (0.3 mM) live imaging buffer (110 mM NaCl, 5.4 mM KCl, 1.2 mM CaCl_2_, 0.8 mM MgCl_2_, 10 mM D-glucose, 10 mM HEPES, pH 7.2). Next, the CNS was immediately transferred into a cell culture dish (Ø 28.2 mm; glass bottom, Ø 20 mm) filled with standard Ca^2+^ (1.2 mM) live imaging buffer. Time-series movies were taken with an inverted Olympus IXplore SpinSR spinning disk confocal super resolution microscope (objective lens, UPLXAPO 40×/0.95 NA, Air) by alternating 488 nm and 561 nm laser channels. The movies were corrected for drifts by Fast4Dreg, and the fluorescence intensities of PEAPODs and mR-GECO1 were quantitated over time by TrackMate in Image J. The dF/F was calculated, and data points were plotted in GraphPad Prism.

### PEAPODs imaging and abundance analysis

The larval CNS was dissected and fixed as described below. After washing off the fixative, samples were incubated in mounting medium (90% glycerol + 10% 1× phosphate buffered saline [PBS]) overnight. Confocal images were taken using an Olympus FV3000 (objective lens, UPLAPO 60×/1.50 NA, Oil). The field of view (FOV) runs 212.1 μm × 212.1 μm, which can accommodate all abdominal segments of the 3^rd^ instar larval VNC. The PEAPODs intensity was determined by measuring the mean intensity of the reconstituted GFP (confocal z-stack projections by maximum intensity) in Image J. The Image J plugin 3D objects counter was used for 3D segmentation, numbering, and volume sizing of PEAPODs. PEAPODs from abdominal segments a1–a2 (cropping, 62.2 μm × 165.3 μm, width × length) are shown in Figs. 2B, 2H, 6A, 6E, and S2A–S2D.

### Thermogenetic activation of DAergic neurons

Control and *th>TrpA1* larvae were cultured in an incubator at 18°C before thermogenetic activation in a 29°C water bath. The larval cohort of each treatment (10-min, 20-min and 30-min) was kept in the water bath until the dissection was completed within the next 10 min.

### Binary expression systems for gene knockdown in astrocytes

We used LexA/lexAop to knock down *pyk* in Figs. 3F, 3G, 4, S3E–S3G, S4D, S4E and S5, while Gal4/UAS was used in the others.

### *pyk* mutagenesis by CRISPR-Cas9

Flies carrying *pyk^TKO.GS^*^03070^ (for the expression of short guide RNA targeting *pyk*) and *nos-Cas9* (for germline Cas9 expression) were crossed to obtain the F1 progeny. Next, F1 males were crossed to a balancer strain, and individual F3 stocks were created by mating single F2 flies to the same balancer strain. Two frameshift deletions of *pyk* (i.e., alleles *pyk*^11^ and *pyk*^7^) were recovered in F3 stocks.

### Antibodies, immunohistochemistry, western blotting, and transmission electron microscopy

The following primary antibodies were used in this study: α-Brp (1:100, nc82, DSHB), α-Repo (1:100, 8D12, DSHB), Alexa Fluor 647 AffiniPure goat anti-HRP (1:200, 123-605-021, Jackson Immunoresearch), α-tubulin (1:5000, ab52866, Abcam), α-Pyk (1:200–1:1000), and α-Nlg2 (1:200). Polyclonal α-Pyk, α-Nlg2 antibodies were produced by immunizing rabbits with synthetic peptides (Pyk, 199 a.a.–212 a.a., EVENGGSLGSRKGV; Nlg2, 295 a.a.–308 a.a., RQRRNTDDTTGEPK) conjugated to keyhole limpet hemocyanin (KLH).

The following secondary antibodies were used in this study: AffiniPure goat anti-mouse Alexa Fluor 647 (1:200, 115-605-062), AffiniPure goat anti-rabbit Alexa Fluor 594 (1:1000, 111-585-045), and peroxidase AffiniPure goat anti-rabbit (1:5000, 111-035-003) (Jackson Immunoresearch).

Immunohistochemistry: Briefly, larval CNSs were dissected in 1× PBS and fixed in 4% PFA for 20 min. After washing off the PFA in 1× PBS, the samples were incubated in 0.3% PBST (1× PBS + 0.3% Triton) for 2 h. Next, the samples were incubated in primary antibody diluted in 0.1% PBST for 72 h at 4°C. The primary antibody was pipetted out, and the samples were washed three times in 0.1% PBST for 10 min each time. The samples were then incubated in secondary antibody diluted in 0.1% PBST for 4 h at room temperature. After removing the secondary antibody, the samples underwent three 10-min washes in 0.1% PBST and one 5-min wash in 1× PBS. Finally, the samples were mounted in Vectashield mounting medium for confocal imaging.

Western blotting: The 2^nd^ instar larval CNS was dissected and homogenized in RIPA (89900, Thermo Fisher) supplemented with proteinase inhibitor cocktail. The lysates were centrifuged (12,000 rpm) at 4°C for 15 min. The supernatants were collected and the total protein concentrations were quantitated using BCA assays. Heat-denatured samples (1 μg/μl) were resolved using 4%–12% gradient SDS-PAGE gels. We followed the standard western blotting procedure for gel-to-membrane (PVDF) protein transfer, membrane blocking, primary antibody incubation (α-Pyk, 1:1000; 4°C, overnight), and secondary antibody binding (peroxidase AffiniPure goat anti-rabbit; 1:5000, room temperature, 1 h) with necessary washing in 1× PBS + 0.1% Tween-20 during each stepwise transition. ECL (34580, Thermo Fisher) was applied on the membrane, and chemiluminescent signals were recorded using a gel imager. Stripping was performed to remove the antibodies from the membrane. The above steps, starting from the primary antibody incubation, were repeated to detect the loading control using α-tubulin.

Transmission electron microscopy: The 3^rd^ instar larval CNS was dissected in 0.1 M PBS and immediately fixed overnight in 2.5% glutaraldehyde (diluted in 0.1 M PBS) at 4 °C. The samples were then rinsed in 0.1 M PBS (three times, 10 min each time) and post-fixed with 2% uranyl acetate for 30 min at room temperature. After a 5-min washing in dH_2_O, the samples were gradually dehydrated in 50%, 70%, 90% (one time,15 min each time) and 100% (twice, 20 min each time) ethanol. The samples were then incubated in 100% acetone (twice, 20 min each time), infiltrated in 50%/50% Epon 812 resin/acetone mixture for 2 h and in 75%/25% Epon 812 resin/acetone mixture overnight. On next day, the samples were carefully orientated in 100% Epon 812 resin in plastic embedding molds and left at 65 °C for polymerization. Ventral nerve cords of the samples were ultrathin sectioned. The sections were placed on copper grids, contrasted in 4% uranyl acetate for 30 min and inspected with a Thermo Scientific Talos L120C TEM at 120 kV.

### FACS, scRNA-seq, data analysis, and cell type annotation

Fifty 3rd instar larval CNSs per genotype were dissected in ice-cold Rinaldini’s solution. The sample collection, cell dissociation, and cell sorting for harvesting mtdTomato^+^ cells by FACS were performed as described by Harzer et al.^2^. The 10× Genomics scRNA-seq was conducted following the manufacturer’s guidelines. Briefly, the 10× library was constructed by amplifying 10× barcoded cDNA obtained by poly (T) RT-PCR from single-cell gel bead-in emulsions (GEMs). For quality control, the insert sizes and the concentration of the 10× library were validated using electrophoresis (Agilent 2100 Bioanalyzer) and qPCR (StepOnePlus Real-Time PCR System), respectively. The 10× library was sequenced on an Illumina HiSeq PE150 sequencer. The RNA sequencing reads were aligned to the released reference *Drosophila_melanogaster*/dmel_r6.54. Data analysis and visualization were performed using Cell Ranger and Seurat. We annotated cell types using customized marker genes from the literature and Garnett software^3–5^.

### Larval tracking and locomotion analysis

The larval locomotion was recorded using frustrated total internal reflection (FTIR)-based imaging (FIM) method at 10 fps for 90 s. For each indicated genotype, 30 larvae were recorded (six larvae per recording × five recordings). The locomotion was analyzed to calculate the sweeping rate, pausing rate, and maximum speed using the FIMTrack built-in tools^6,7^. The bending strength (i.e., median bending angle), peristalsis efficiency (i.e., distance per peristalsis), and peristalsis frequency (i.e., peristalses per second) were quantitated by pyFIM (Python package for analysis of FIMTrack data, https://github.com/schlegelp/pyfim). The minimum track length in frames was set to 50. Data filtering owing to larval collision, larval disappearance, and return in FOV may result in slightly more or less than 30 data points per genotype depending on the analysis performed. The sweeping rate was calculated as follows: the number of frames the larva sweeps/the total number of frames the larva appears × 100%. The pausing rate was calculated as follows: the number of frames the larva pauses/the total number of frames the larva appears × 100%.

### Statistical analysis

Sample sizes were chosen to meet published standards. All experiments were technically repeated at least twice (see figure legends for exact number) with similar results. Regarding single-cell RNA sequencing, dataset integration was performed to correct for technical differences between the control and astrocyte *pyk* knockdown. All data were presented as the mean ± SEM. The *P*-value was obtained by performing one-way ANOVA analysis with Tukey’s test, unpaired t-test, or Kolmogorov-Smirnov test in GraphPad Prism, as indicated in the figures and figure legends.

**Table S1.**
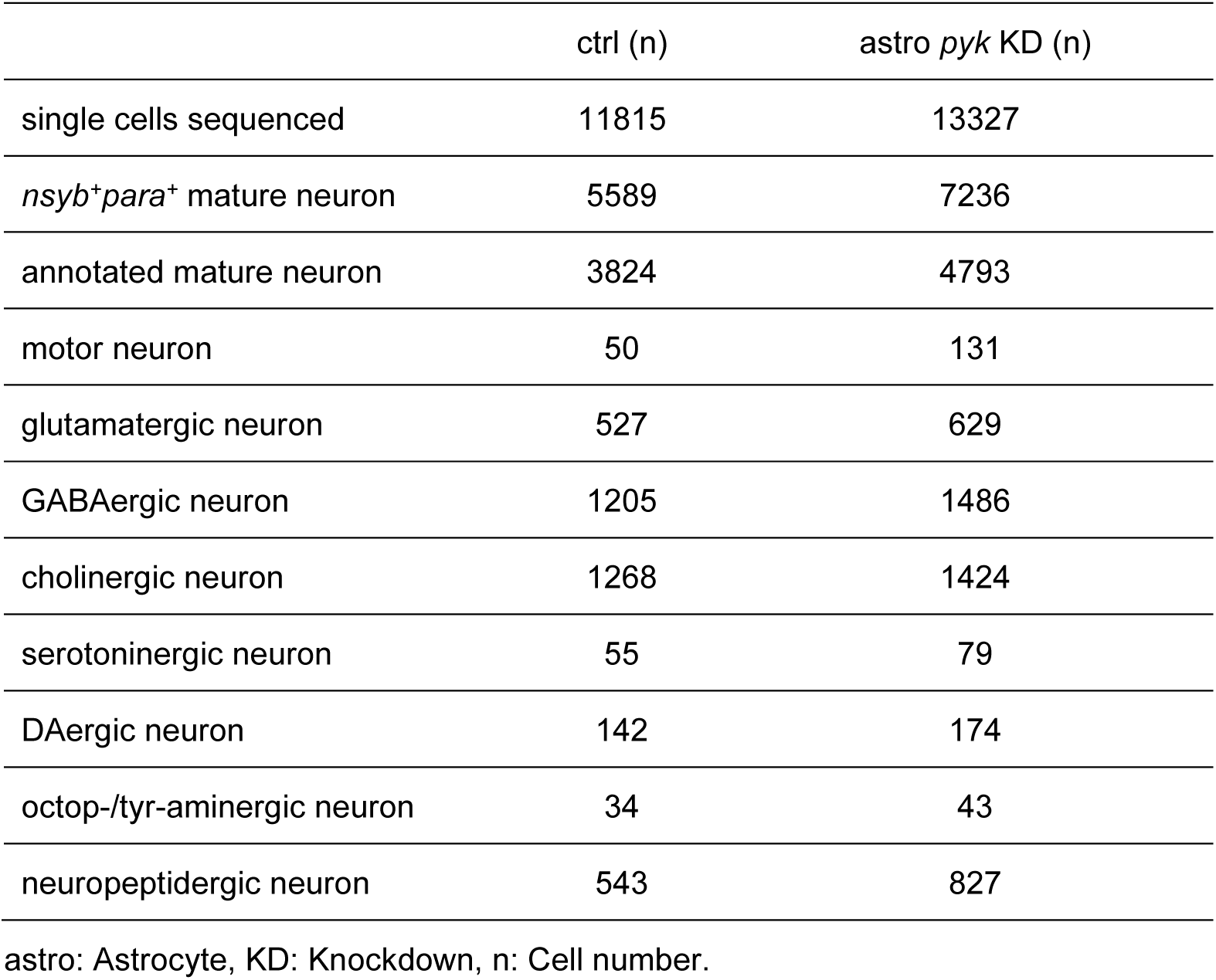
The summary list of cells identified by sc-RNA seq and cell type annotation

|  | ctrl (n) | astro *pyk* KD (n) |
| --- | --- | --- |
| single cells sequenced | 11815 | 13327 |
| *nsyb<sup>+</sup>para<sup>+</sup>* mature neuron | 5589 | 7236 |
| annotated mature neuron | 3824 | 4793 |
| motor neuron | 50 | 131 |
| glutamatergic neuron | 527 | 629 |
| GABAergic neuron | 1205 | 1486 |
| cholinergic neuron | 1268 | 1424 |
| serotonergic neuron | 55 | 79 |
| DAergic neuron | 142 | 174 |
| octop-/tyr-aminergic neuron | 34 | 43 |
| neuropeptidergic neuron | 543 | 827 |

astro: Astrocyte, KD: Knockdown, n: Cell number.

**Table S2.**
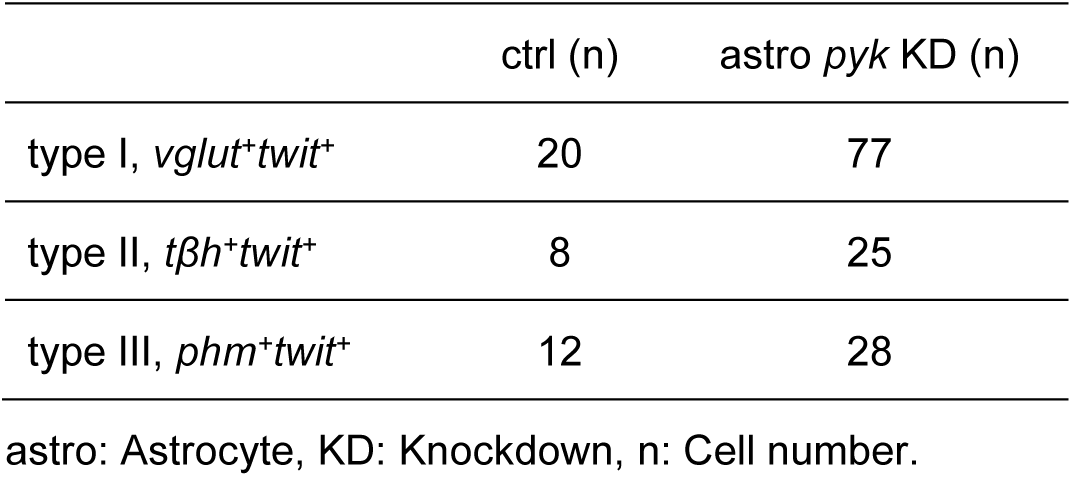
Motor neuron subclasses connected to *Drosophila* larval DAergic system

|  | ctrl (n) | astro *pyk* KD (n) |
| --- | --- | --- |
| type I, *vglut<sup>+</sup>twit<sup>+</sup>* | 20 | 77 |
| type II, *tβh<sup>+</sup>twit<sup>+</sup>* | 8 | 25 |
| type III, *phm<sup>+</sup>twit<sup>+</sup>* | 12 | 28 |

astro: Astrocyte, KD: Knockdown, n: Cell number.

**Table S3.**
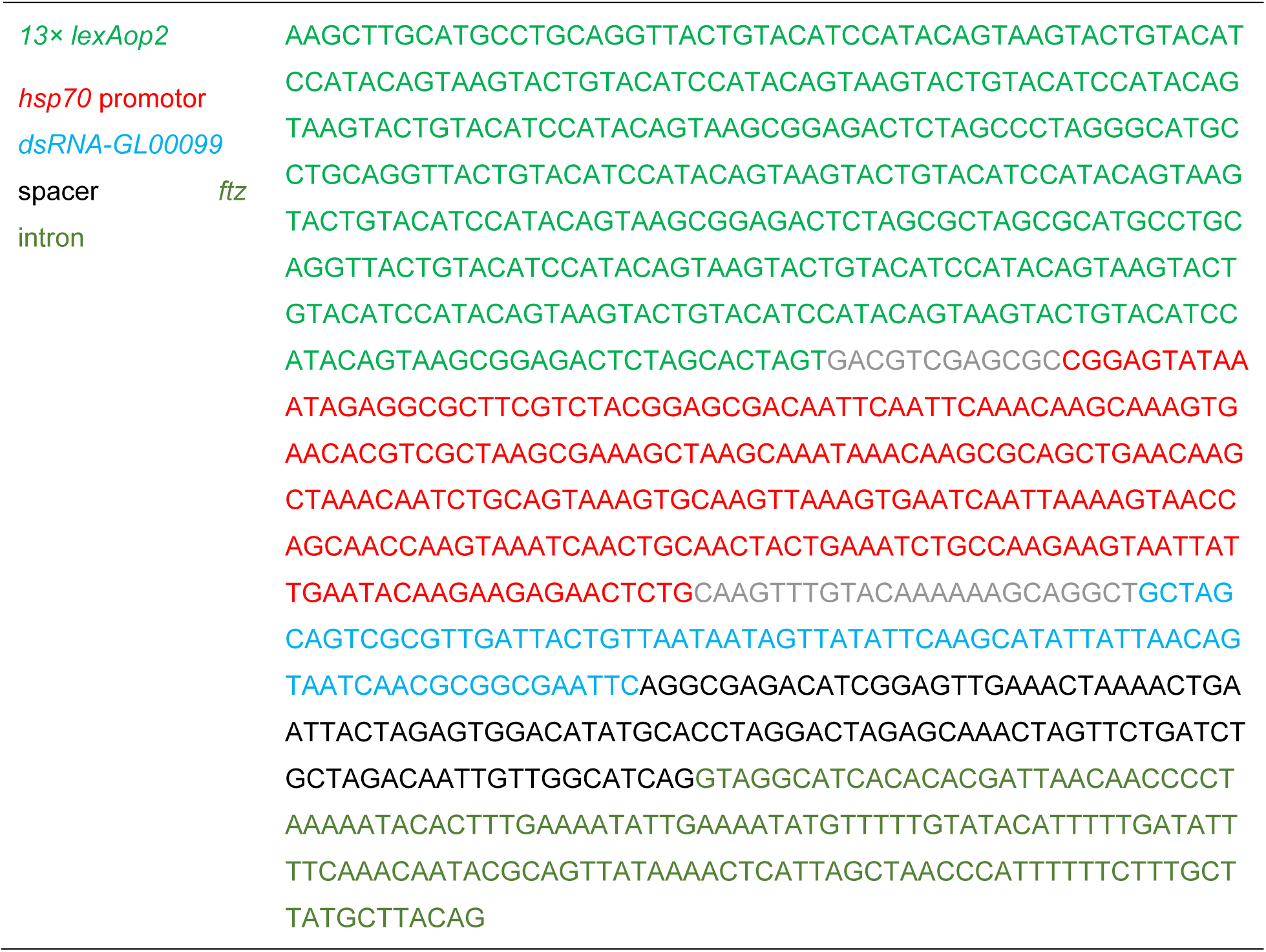
The DNA sequence assembled for creating *lexAop2-pyk^RNAi^* construct

**Table S4.** Stock list

| Name | Stock Number | Stock Center/Reference |
| --- | --- | --- |
| *UAS-pfk<sup>RNAi#1</sup>* | 34336 | BDSC |
| *UAS-pfk<sup>RNAi#2</sup>* | TH201500692.S | THFC |
| *UAS-eno<sup>RNAi#1</sup>* | 26300 | BDSC |
| *UAS-eno<sup>RNAi#2</sup>* | TH02508.N | THFC |
| *UAS-pyk<sup>RNAi#1</sup>* | 35218 | BDSC |
| *UAS-pyk<sup>RNAi#2</sup>* | TH03685.N | THFC |
| *lexAop2-pyk<sup>RNAi</sup>* | N/A | this study |
| *UAS-nlg2<sup>RNAi</sup>* | 28331 | BDSC |
| *UAS-nrx1<sup>RNAi</sup>* | 27502 | BDSC |
| *UAS-brp<sup>RNAi#1</sup>* | 80449 | BDSC |
| *UAS-brp<sup>RNAi#2</sup>* | 25891 | BDSC |
| *w<sup>1118</sup>* | 5905 | BDSC |
| *10× UAS-IVS-mtdTomato* | 32222 | BDSC |
| *th-Gal4* | 8848 | BDSC |
| *th-LexA* | 99051 | BDSC |
| *UAS-TrpA1* | 26263 | BDSC |
| *pyk<sup>TKO.GS03070</sup>* | 78770 | BDSC |
| *nos-Cas9* | TH00788.N | THFC |
| *Brp-FSF-GFP* | 55753 | BDSC |
| *trans-Tango QUAS-mtdTomato-3× HA* | 90924 | BDSC |
| *hs-FLPG5* | 58356 | BDSC |
| *20× UAS-FLPG5* | 55806 | BDSC |
| *UAS-GFP-KDEL* | 9898 | BDSC |
| *UAS-mKcnj2* | 600376 | BDSC |
| *UAS-mKcnj2.nc* | 600377 | BDSC |
| *UAS-TNT-G* | 28838 | BDSC |
| *UAS-TNT-Q* | 28839 | BDSC |
| *tub>Gal80>* | 38881 | BDSC |
| *UAS-CD4-spGFP<sup>1-10</sup>* | 93017 | BDSC |
| *lexAop-CD4-spGFP<sup>11</sup>* | 93018 | BDSC |
| *alrm-LexA* |  | Ma et al.<sup>1</sup> |
| *alrm-Gal4* |  | Doherty et al.<sup>8</sup> |
| *UAS-brp.S-mStraw* |  | Fouquet et al.<sup>9</sup> |
| *UAS-mR-GECO1* |  | Ma et al.<sup>10</sup> |
| *repo-FLP* |  | Stork et al.<sup>11</sup> |
| *UAS-mCD8-GFP* |  | Lee & Luo<sup>12</sup> |

BDSC: Bloomington *Drosophila* Stock Center, THFC: TsingHua Fly Center.

**Fig. S1.**
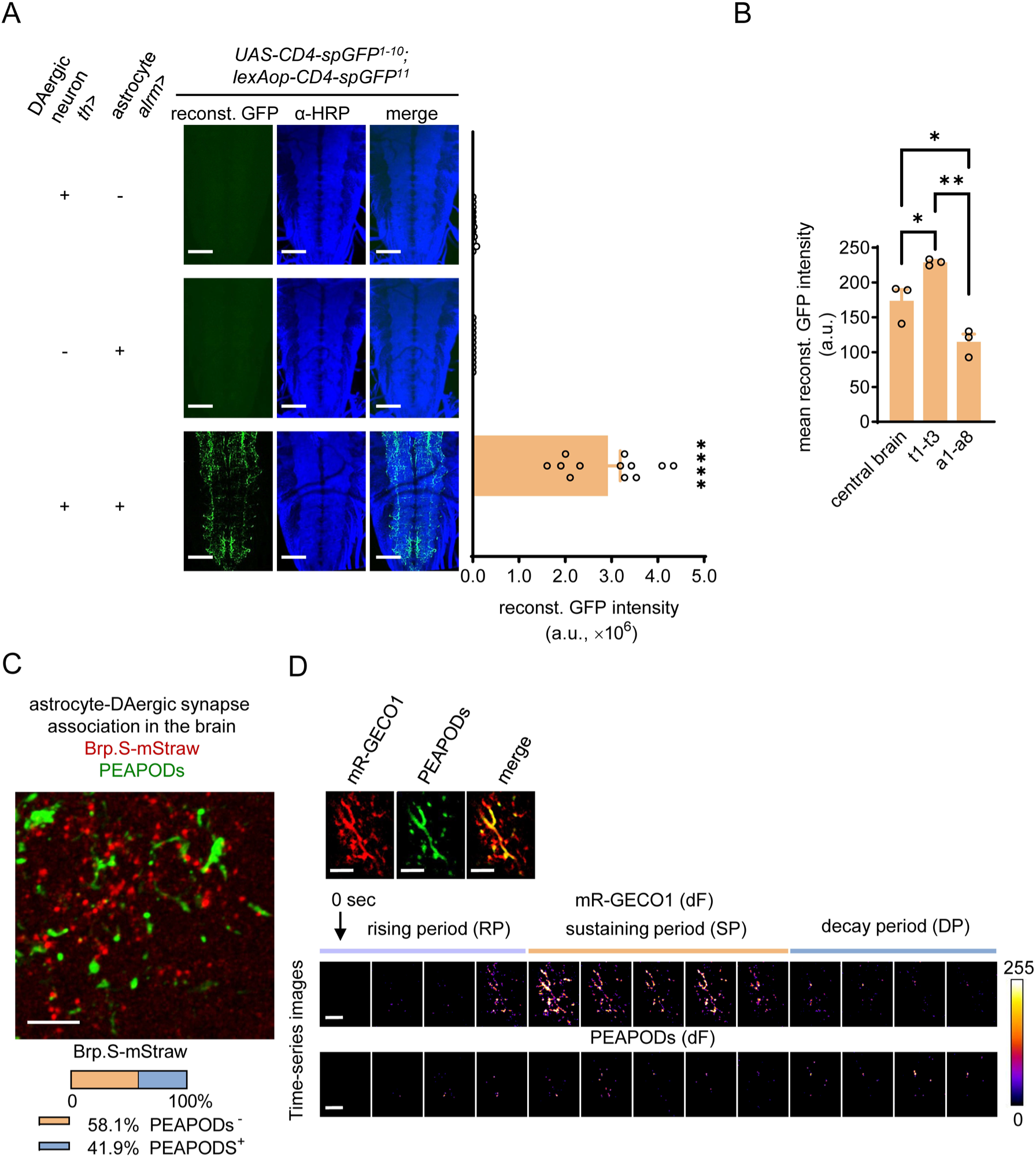
Trans astrocyte-DAergic neuronal GFP reconstitution, and PEAPODs remodeling during spontaneous DAergic neuronal activation. (A) Trans astrocyte-DAergic neuronal reconstituted GFP (reconst. GFP) (n = 12) (scale bars: 50 µm). α-HRP (horseradish peroxidase) stains neural tissues. The drivers *th-Gal4* (*th*) and *alrm-LexA* (*alrm*) were used here. a.u.: Arbitrary unit. (B) Regional variations in reconstituted GFP (reconst. GFP) intensity across the larval ventral nerve cord (VNC; thoracic segments, t1–t3; abdominal segments, a1–a8) and central brain (n = 3). a.u.: Arbitrary unit. (C) Single-cell PEAPODs and synaptic labeling of the DAergic neuron in larval brain (n = 10) (scale bars: 5 µm). (D) Dual-color live imaging of PEAPODs and spontaneous DAergic neuronal activation. Active DAergic neuronal projections (labeled by mR-GECO1) and their associated PEAPODs. Time-series dF (Ft–F_0_) images show the differential fluorescence intensity at the time point of t second as compared to the baseline (0 s) (scale bars: 10 µm). Imaging time interval, 0.7 s. One-way ANOVA with Tukey’s test. \**P* < 0.05, \*\**P* < 0.01, \*\*\*\**P* < 0.0001. FC: Fold change.

**Fig. S2.**
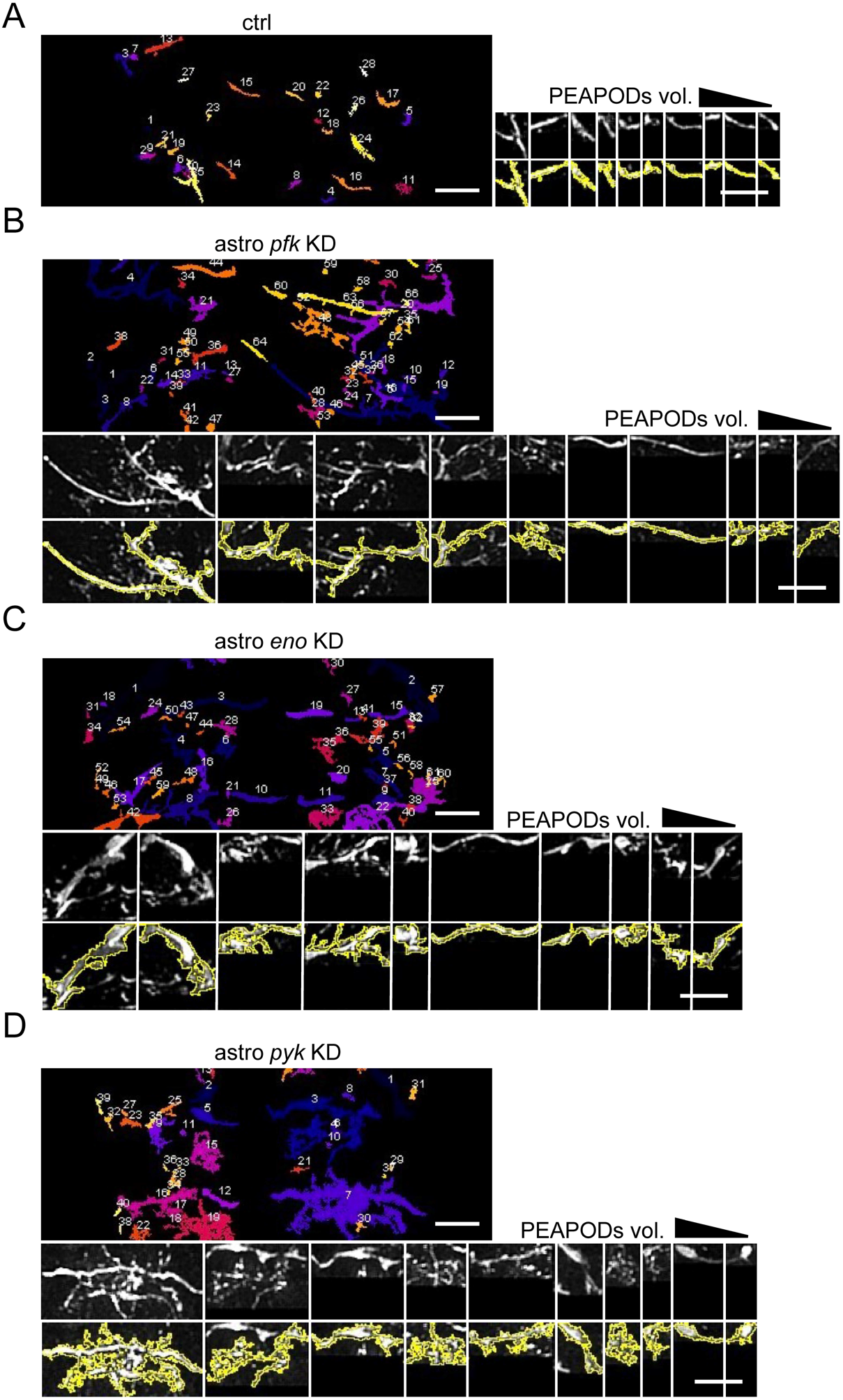
The knockdown of glycolytic genes that stimulate PEAPODs. (A–D) Three-dimensional segmentation and numbering of individual PEAPODs (scale bars: 25 µm). Ten largest PEAPODs from the indicated genotypes were sorted by volume (vol.), and they are delineated by the orange line (scale bars: 25 µm). astro: Astrocyte, KD: Knockdown.

**Fig. S3.**
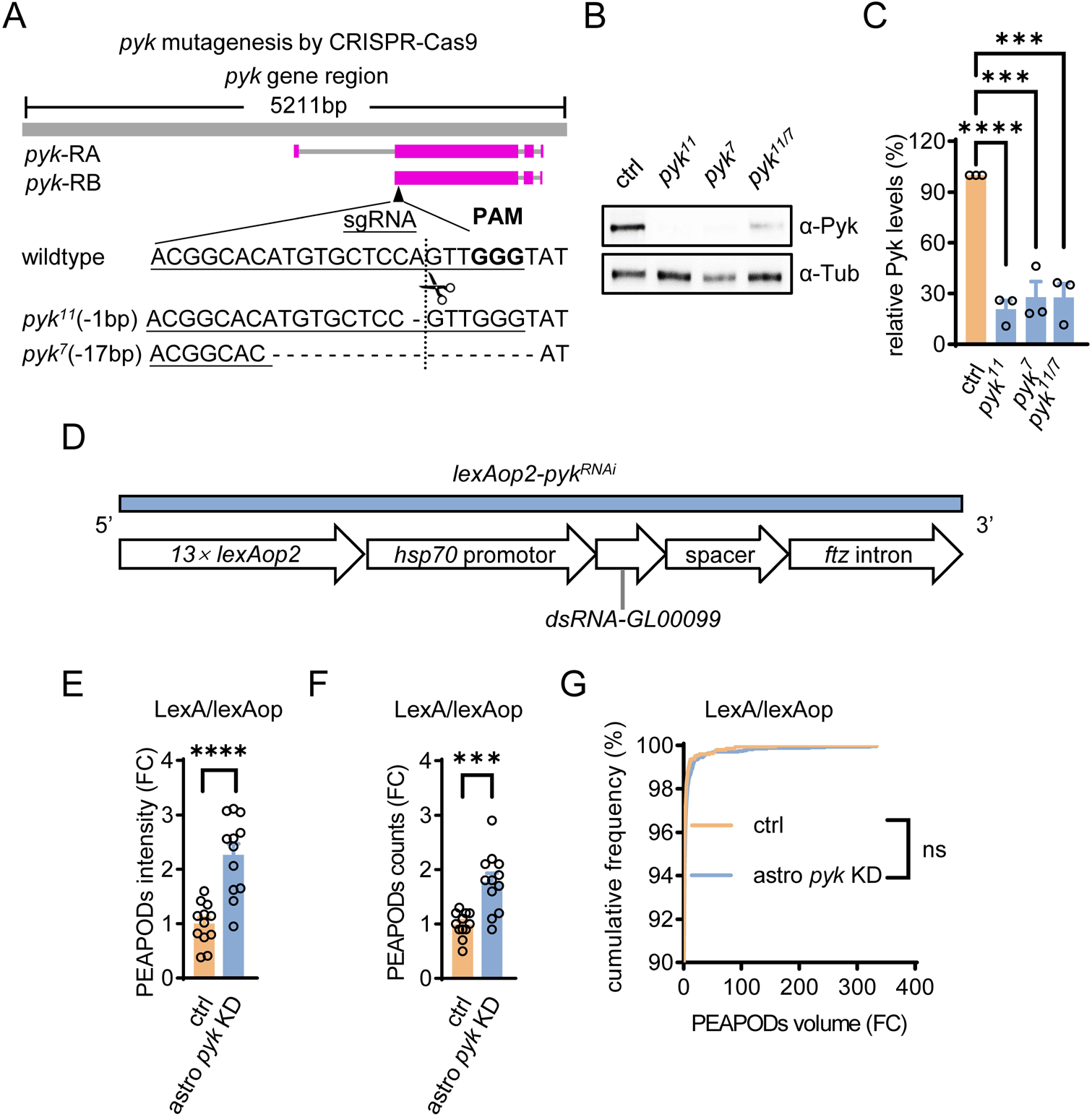
*pyk* mutagenesis using CRISPR-Cas9, *lexAop2-pykRNAi* construct assembly. (A) Schematics of the *pyk* gene region, two predicted transcripts (*pyk*-RA, -RB), short guide RNA (sgRNA) for guiding Cas9 to the *pyk* locus, and DNA lesions of the two mutated alleles, *pyk11* and *pyk7*. PAM: Protospacer adjacent motif. (B and C) Western blotting using α-Pyk antibody (B), and quantitated relative Pyk protein levels (C). Loading control, α-Tub (tubulin). Genotypes as indicated (n = 3). (D) Schematic illustration of *lexAop2-pykRNAi* construct assembly. *dsRNA-GL00099* is the TRiP RNAi amplicon (used in *pyk* RNAi line #1) for generating short hairpin RNA (shRNA) targeting *pyk*. The *ftz* intron was purposed to facilitate shRNA nuclear export. The illustrated elements (DNA sequence included in Table S3) arranged as such were cloned into the *pUASTattB* backbone. Note that *5× UAS* was replaced by 13 copies of *lexAop2* in tandem (*13× lexAop2*). (E–G) PEAPODs abundance analysis. Astrocyte (astro) *pyk* knockdown (KD) using LexA/lexAop system. Genotypes as indicated (n = 12; in G, 1970–3519 individual PEAPODs in total). (C), one-way ANOVA with Tukey’s test. (E), (F), unpaired t-test. (G), Kolmogorov-Smirnov test. \*\*\**P* < 0.001, \*\*\*\**P* < 0.0001. ns: Not significant, FC: Fold change.

**Fig. S4.**
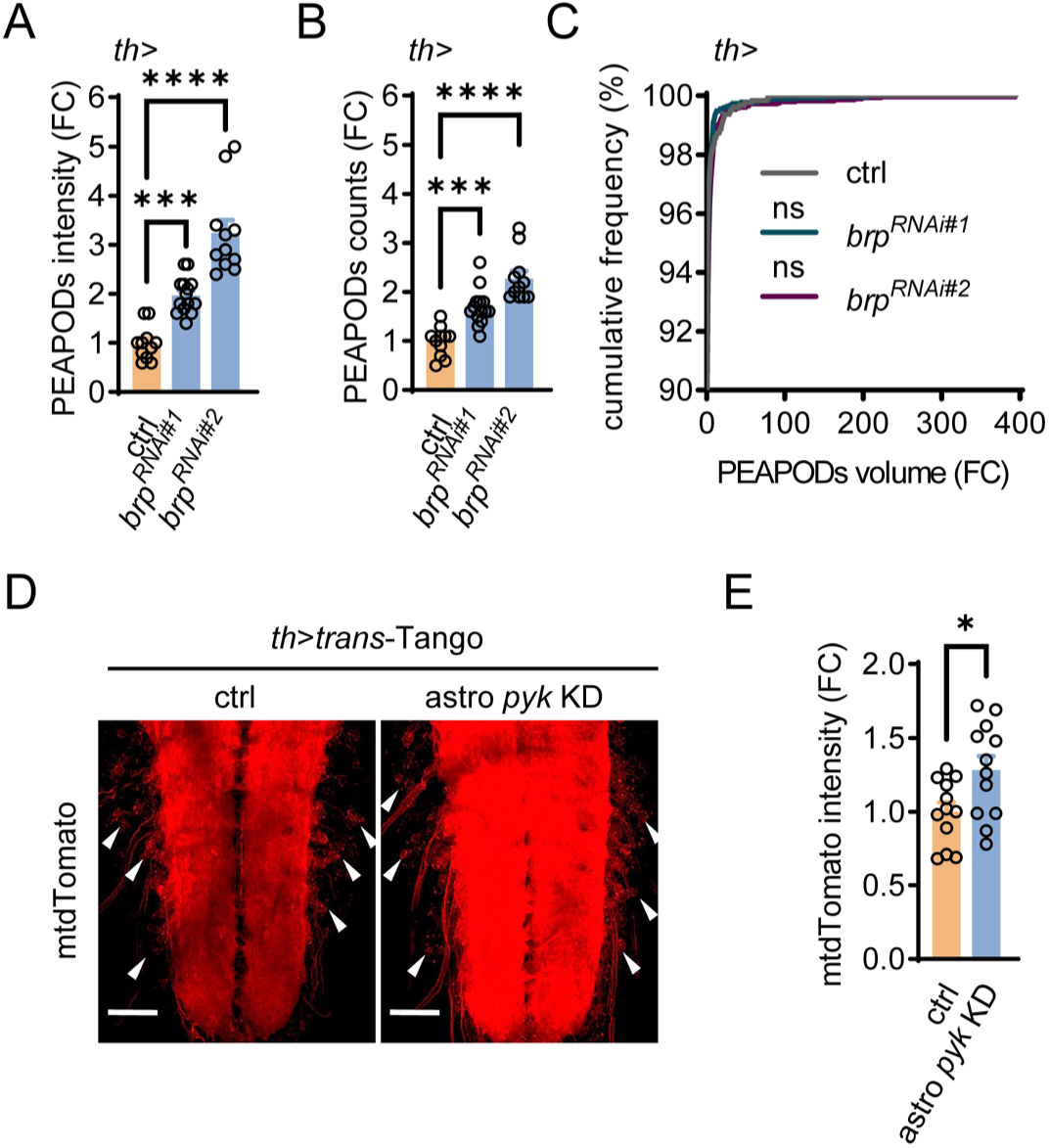
PEAPODs in DAergic neuron *brp* knockdown (KD) transsynaptic (DAergic synapse) tracing by *trans*-Tango. **(A–C)** PEAPODs abundance analysis. Genotypes as indicated (n = 11–14; in C, 1267–2876 individual PEAPODs in total). *th*: DAergic neuron driver. (D and E) mtdTomato intensity. The postsynaptic partners of DAergic neurons express myristoylated tdTomato (mtdTomato) (scale bars: 50 µm). Arrowheads denote the neuron somata. Genotypes as indicated (n = 12). *trans*-Tango: Transsynaptic labeling system, astro: Astrocyte, KD: Knockdown. (A), (B), one-way ANOVA with Tukey’s test. (C), Kolmogorov-Smirnov test. (E), unpaired t-test. \**P* < 0.05, \*\*\**P* < 0.001, \*\*\*\**P* < 0.0001. ns: Not significant, FC: Fold change.

**Fig. S5.**
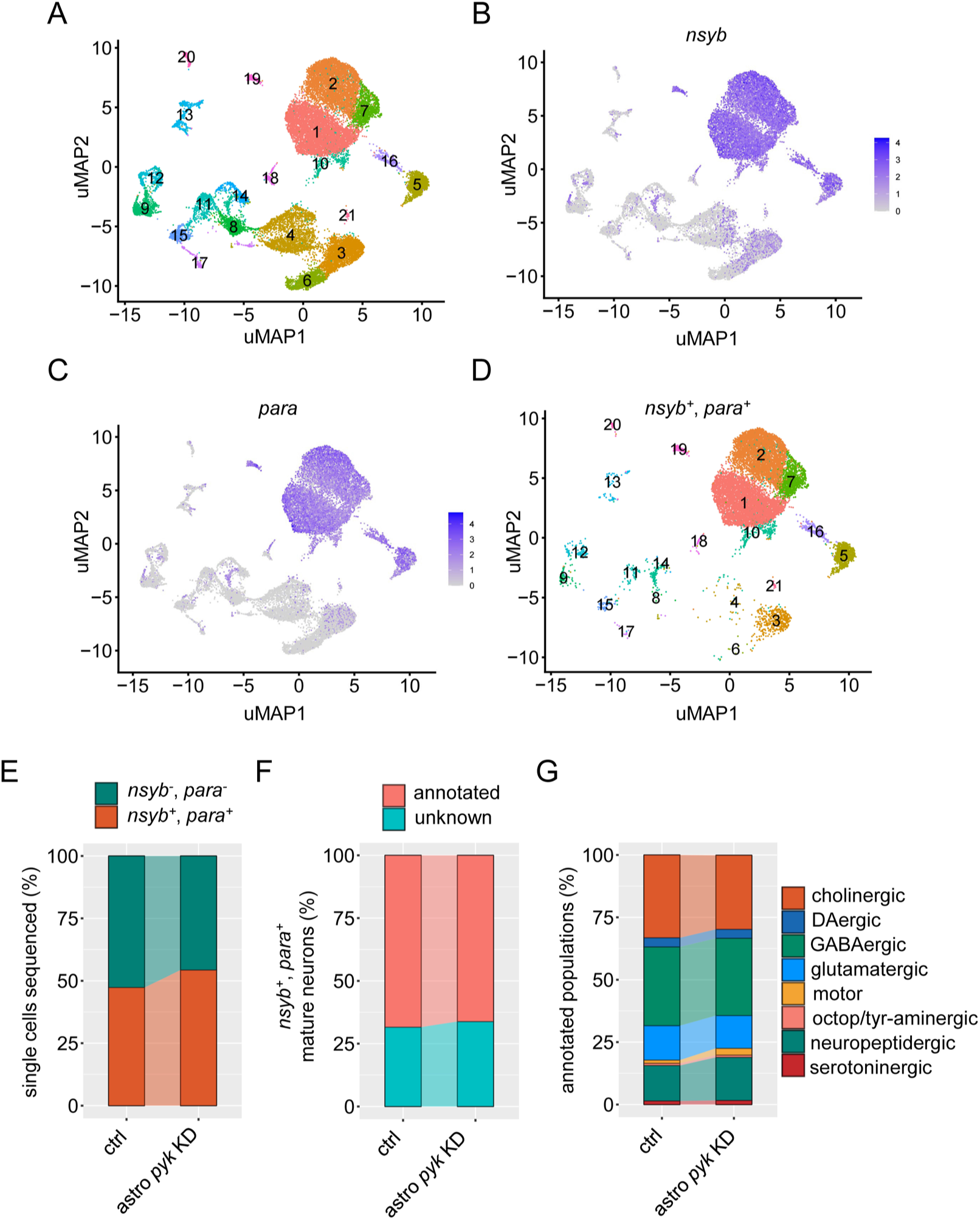
scRNA-seq and cell type annotation of postsynaptic partners connected to DAergic neurons. (A) Integrated clustering of all cells sequenced from ctrl (n = 11815) and astrocyte (astro) *pyk* knockdown (KD) (n = 13327) in uMAP. uMAP: Uniform manifold approximation and projection. (B and C) Single cells expressing *nsyb* (B), *para* (C). Note that the cells showed a tendency to converge in the upper right groups, e.g., clusters 1, 2, 5, 7, 10, 16, and 19. Color scale: Normalized gene expression level. (D) Highlighted *nsyb+para+* cells in clusters. (E) Percentages of *nsyb-para-* cells and *nsyb+para+* mature neurons in all sequenced cells. (F) Percentages of (un)annotated neurons in mature neurons. (G) Percentages of defined populations in annotated mature neurons.

**Figure S6.**
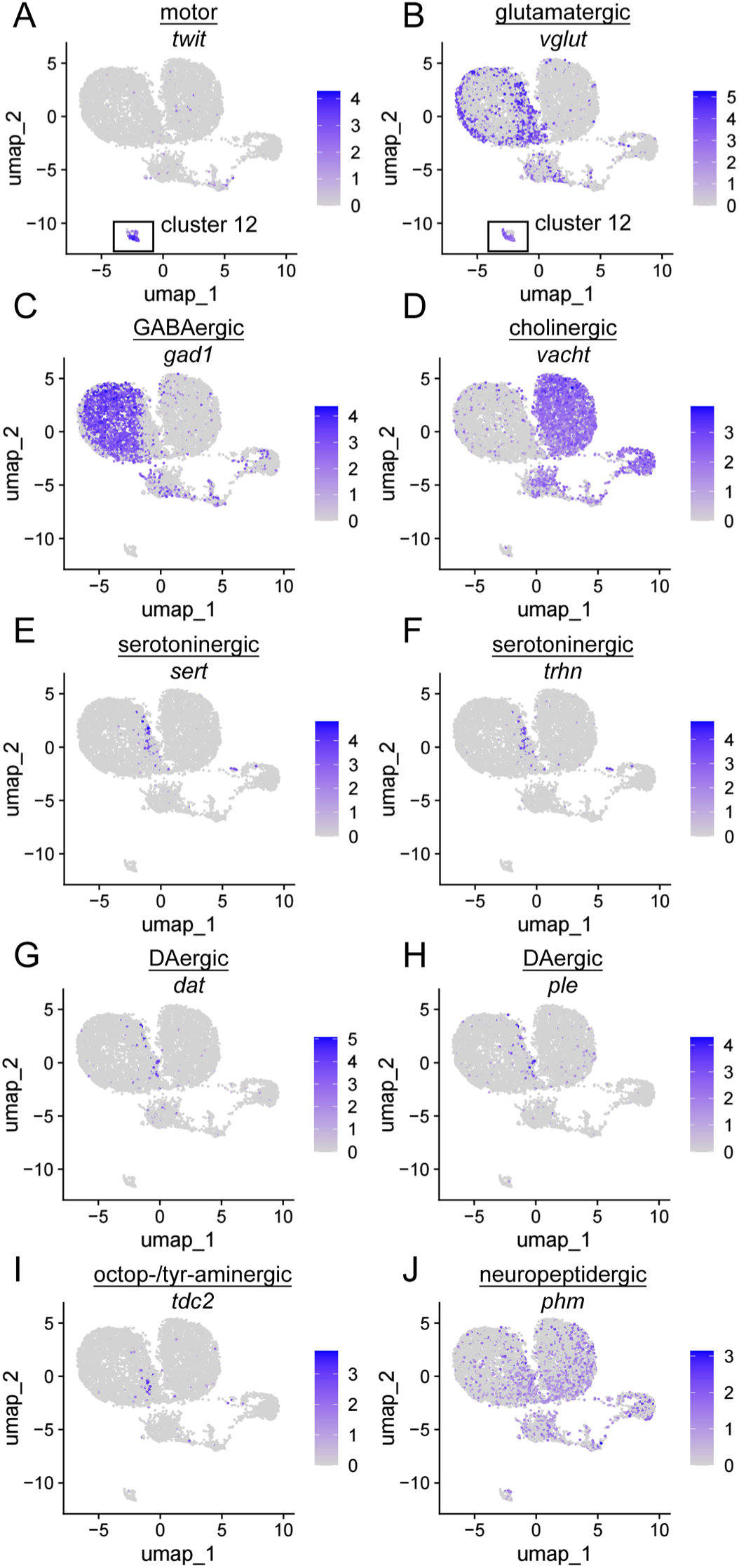
Annotated populations defined by unique marker genes. (A and B) Motor neuron (MN) and marker gene *twit* (A). Glutamatergic neuron and marker gene *vglut* (B). Note that MNs gathered in cluster 12, which was also *vglut*+. (C) GABAergic neuron and marker gene *gad1*. (D) Cholinergic neuron and marker gene *vacht*. (E and F) Serotoninergic neuron and marker genes: *sert* (E), *trhn* (F). (G and H) DAergic neuron and marker genes: *dat* (G), *ple* (H). (I) Octop-/tyr-aminergic neuron and marker gene: *tdc2*. (J) Neuropeptidergic neuron and marker gene: *phm*. Integrated clustering contains all annotated cells from the ctrl and astrocyte *pyk* knockdown. uMAP: Uniform manifold approximation and projection, color scale: Normalized gene expression level.

**Movie S1 (separate file).** Z-axis confocal sections showing DAergic neuronal branches (red), PEAPODs (green) and the presynaptic marker Brp (magenta)

**Movie S2 (separate file).** Dual-color live imaging of spontaneous DAergic neuronal activation (shown by mR-GECO1) and PEAPODs

