## Supplementary figures and images for "Metabolic basis of the astrocyte-synapse interaction governs dopaminergic-motor connection"

### Supplemental video S1

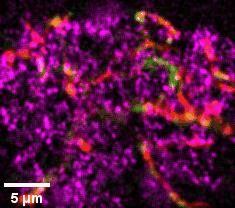

### Supplemental video S2

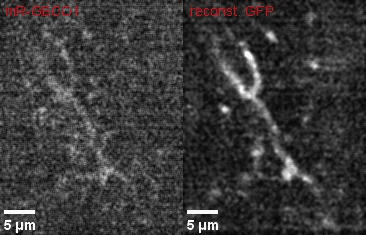
